# Combined Mek inhibition and *Pparg* activation Eradicates Muscle Invasive Bladder cancer in a Mouse Model of BBN-induced Carcinogenesis

**DOI:** 10.1101/2023.08.19.553961

**Authors:** Tiffany Tate, Sakina A Plumber, Hikmat Al-Ahmadie, Xiao Chen, Woonyoung Choi, Chao Lu, Aaron Viny, Ekatherina Batourina, Krisjian Gartensson, Besmira Alija, Andrei Molotkov, Gregory Wiessner, James McKiernan, David McConkey, Colin Dinney, Bogdan Czerniak, Cathy Lee Mendelsohn

## Abstract

Bladder cancers (BCs) can be divided into 2 major subgroups displaying distinct clinical behaviors and mutational profiles: basal/squamous (BASQ) tumors that tend to be muscle invasive, and luminal/papillary (LP) tumors that are exophytic and tend to be non-invasive. *Pparg* is a likely driver of LP BC and has been suggested to act as a tumor suppressor in BASQ tumors, where it is likely suppressed by MEK-dependent phosphorylation. Here we tested the effects of rosiglitazone, a *Pparg* agonist, in a mouse model of BBN-induced muscle invasive BC. Rosiglitazone activated *Pparg* signaling in suprabasal epithelial layers of tumors but not in basal-most layers containing highly proliferative invasive cells, reducing proliferation but not affecting tumor survival. Addition of trametinib, a MEK inhibitor, induced *Pparg* signaling throughout all tumor layers, and eradicated 91% of tumors within 7-days of treatment. The 2-drug combination also activated a luminal differentiation program, reversing squamous metaplasia in the urothelium of tumor-bearing mice. Paired ATAC-RNA-seq analysis revealed that tumor apoptosis was most likely linked to down-regulation of Bcl-2 and other pro-survival genes, while the shift from BASQ to luminal differentiation was associated with activation of the retinoic acid pathway and upregulation of Kdm6a, a lysine demethylase that facilitates retinoid-signaling. Our data suggest that rosiglitazone, trametinib, and retinoids, which are all FDA approved, may be clinically active in BASQ tumors in patients. That muscle invasive tumors are populated by basal and suprabasal cell types with different responsiveness to *PPARG* agonists will be an important consideration when designing new treatments.

## Introduction

Urothelial carcinoma of the bladder (bladder cancer; BC) is currently the 6th most common form of cancer in the United States with about 81,000 new cases and 17,000 deaths per year. It occurs 3 times more frequently in men than in women, but women tend to have more aggressive disease ^1^. Most cases of BC (70%) present as non-muscle invasive bladder cancer (NMIBC) that includes patients with stage Ta and T1 tumors as well as carcinoma in situ. All NIMBCs are treated with complete transurethral resection, and patients with high-risk disease are also treated with adjuvant intravesical BCG. However, 50-70% of patients experience recurrence and up to 20% of high-grade NMIBCs progress to muscle invasive bladder cancer [MIBC; ^2^]. MIBC is associated with a 50% five-year survival rate, and metastatic progression almost always leads to death ^3,4^. Neoadjuvant cisplatin-based chemotherapy followed by definitive surgery (radical cystectomy) has been the preferred treatment for MIBC since the early 2000s (Adamo et al., 2005; Als et al., 2008; Dash et al., 2008; Dogliotti et al., 2007), although radiation-based trimodal therapy (TMT) is equally effective in well-selected patients ^5^. Immune-checkpoint inhibitors (ICIs) such as atezolizumab and pembrolizumab are active in approximately 20% of patients with advanced and metastatic disease^6–8^, and novel combinations of ICIs and antibody-drug conjugates are displaying even stronger activity, although the durability of this benefit is still unclear. Overall, there is still a great need to identify new avenues for treatment of BC.

Bladder cancers are clinically and biologically heterogeneous, and genomic profiling studies performed over the last decade have identified some of the underlying mechanisms. At the highest level they can be subdivided into two subtypes based on their differential expression of biomarkers that are expressed within the different cell layers that are present in the normal urothelium. Almost all NMIBCs and about half of MIBCs express a similar set of markers as are observed in intermediate and superficial cells in the healthy urothelium (hereafter referred to as uro/luminal markers), including *PPARG*, *FOXA1* and *GATA3*. The other half of MIBCs express a set of markers shared with basal cells in the normal urothelium and squamous cells, many of which are not detected in the healthy urothelium (Cancer Genome Atlas Research, 2014; Choi et al., 2014; Damrauer et al., 2014; McConkey et al., 2015a; McConkey et al., 2015b; Robertson et al., 2017; Saito et al., 2018; Sjodahl et al., 2012; Warrick et al., 2016). The most recent consensus classification identified 6 molecular subtypes of BC with distinct mutational landscapes and distinct responses to therapies: luminal papillary, luminal non-specified, luminal unstable, stroma-rich, neuroendocrine and basal/squamous ^2^. Bladder cancer subtypes also display distinct immune signatures: BASQ tumors are highly immune infiltrated and appear to be more responsive to ICIs, whereas LP tumors are poorly infiltrated with immune cells and hence less likely to respond to immune checkpoint inhibitors ^2,7,9,10^.

*Pparg* is a critical regulator of urothelial differentiation both in mice and in humans and is also important for specification of the bladder cancer subtype, promoting luminal differentiation and opposing BASQ differentiation ^11–15^. *Pparg* belongs to the nuclear receptor superfamily of ligand activated transcription factors. It forms heterodimers with *Rxr*, another nuclear receptor family member, that bind to response elements located in regulatory sequences of target genes ^16^. Its ligands include dietary lipids and eicosanoids as well as synthetic agonists such as thiazolidinediones that are used clinically to manage type 2 diabetes. In the absence of ligand, *Pparg*/Rxr heterodimers are in a repressive conformation associated with co-repressors including SMRT and NCoR and HDAC ^17^. Ligand binding induces a conformational change in the complex, promoting the release of co-repressors and recruitment of co-activators that include Med1 and NCoa family members.

Recent studies in “mesenchymal” breast cancer mouse models indicated that while the thiazolidinedione rosiglitazone had some anti-tumor activity, it is much more potent when it was combined with the MEK inhibitor trametinib^18^. These observations prompted us to determine whether a similar protocol could be efficacious in the BBN mouse model of BASQ MIBC. Here we show that treatment with rosiglitazone (Rosi) alone slowed tumor growth and decreased proliferation. Interestingly, Rosi induced *Pparg* expression and transcirptional activity only in suprabasal layers of Ba/Sq tumors, whereas treatment with a combination of Rosi plus trametinib (Tram) induced *Pparg* signaling throughout tumors, resulting in widespread apoptosis within 7 days of treatment and reducing tumor volumes by 91%. The 2-drug combination also reversed squamous metaplasia and restored normal differentiation in the urothelium of tumor-bearing mice. Paired ATAC-RNA-seq analysis revealed that tumor cell apoptosis was linked to down-regulation of Bcl2 and other pro-survival genes, while the shift from BASQ to luminal differentiation was likely driven by retinoid signaling, which was activated down-stream of *Pparg*. In addition, the 2-drug combination induced expression of *Kdm6a*, a lysine demethylase that facilitates retinoid-signaling in urothelial cells and in mouse organoids ^19^. Our data suggest that Rosi, Tram, and retinoids, including Tretinoin, which are FDA approved, may be efficacious in treating MIBC in a clinical setting. That muscle invasive tumors are populated by basal and suprabasal cell types with different drug sensitives will be an important consideration when designing treatments for MIBC.

## RESULTS

### Combined Rosiglitazone and Trametinib induce tumor regression in a mouse model of BBN-induced MIBC

*Pparg*, which is undetectable in in BASQ tumors at the level of mRNA and protein, has been proposed to be a potential tumor suppressor in MIBCs ^20,21^. To directly test this possibility, we tested the effects of the synthetic agonist rosiglitazone (hereafter referred to as Rosi), on BASQ tumor growth and/or survival. We used a mouse model of BBN [N-butyl-n-(4-hydroxybutyl)-nitrosamine]-induced carcinogenesis that produces BASQ tumors after 4-5 months of treatment ^22–24^. Wild type mice were exposed to BBN for 5 months then treated with Rosi or vehicle by oral daily gavage for one month. Ultrasound analyses of tumor growth in control animals treated with vehicle revealed robust growth where tumors size increased on average by 3560% (Table 1; Fig. 1A,B,I), whereas in Rosi treated mice, tumor growth was markedly reduced (Table 1; Fig. 1C,D,I). These observations indicate that Rosi slowed growth but was not sufficient to eradicate the BBN-induced BASQ tumors.

**Figure 1.**
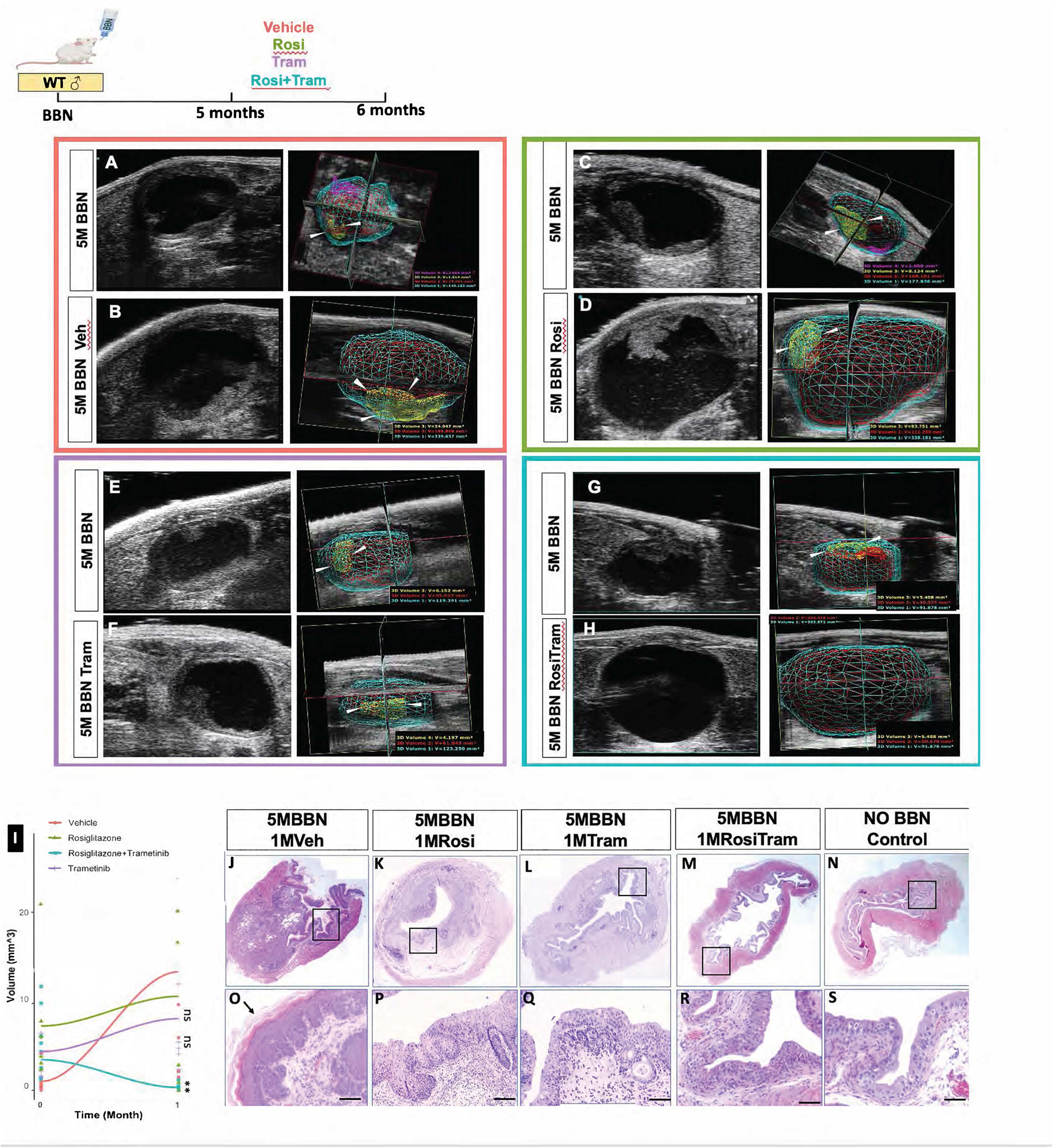
Combined Rosi+Tram treatment reduces tumor burden in mice with BBN-induced MIBC. Wildtype male mice were given BBN (0.05%) in drinking water *ad libidum* for 5 months. Mice were then treated for one month by daily via oral gavage with vehicle (Control), Rosi alone, Tram alone, or with a combination of Rosi and Tram. Tumor volume was measured prior to euthanasia, and bladders were collected. Ultrasound images and 3D reconstruction of bladders were obtained from mice before and after BBN exposure. **A,C**. Representative images of ultrasound imaging of a mouse exposed to BBN for 5 months before (**A)** and after **(B)** administration of Vehicle for 1 month. Ultrasound image of a mouse exposed to BBN for 5 months before (**C)** and after **(D)** administration of Rosi for 1 month. Ultrasound image of a mouse exposed to BBN for 5 months before (**E)** and after **(F)** 1 month of treatment with Rosi. Ultrasound images and 3D reconstruction of a bladder from a mouse treated for 5 months with BBN before **(E)** or after **(F)** treatment for 1 month with Tram. Ultrasound images and 3D reconstruction of bladders from mice treated for 5 months with BBN before **(G)** and after **(H)** 1 month of combined Rosi/Tram treatment. **(I).** Tumor volume measurements based on 3D reconstruction before and after 1M vehicle, 1M Rosi, 1M Tram or 1M Rosi+Tram. Significance calculated by two way Anova **p=0.0021. **J-N.** Panoramas showing histological analysis of a bladders from BBN exposed mouse after 1 month of vehicle **(J),** after 1 month of Rosi treatment (**K**), after 1 month of Tram (**L**), after 1 month of Rosi+Tram (**M**) and untreated wild type control **(N). (O-S)** Panoramas showing histology of samples after treatment with vehicle **(J)**, Rosi **(K)**, Tram **(L)** Rosi+Tram (M) untreated adult mouse control **(N). (O-S).** Magnified regions in boxed regions shown in images J-N. In images of showing 3D bladder reconstruction, blue denotes outer muscle layer, red denotes urothelium, yellow and pink denote tumors. Arrowhead in (O) denotes keratinized squamous differentiation in a BBN-induced tumor. Scale bars: 50μm.

**Table 1.** Tumor volume based on ultrasound from before and after treatments.

| Treatment | Number | Before Volume (mm<sup>3</sup>) | After Volume (mm<sup>3</sup>) | Change (mm<sup>3</sup>) | Average Change (mm<sup>3</sup>) | Change (%) | Average Change (%) |
| --- | --- | --- | --- | --- | --- | --- | --- |
| 5M BBN<br>1M Vehicle | 1 | 0.265 | 2.190 | 1.925 | 12.604 | 726.4% | 35360.5% |
|  | 2 | 0.729 | 6.057 | 5.328 |  | 730.9% |  |
|  | 4 | 4.278 | 24.047 | 19.769 |  | 462.1% |  |
|  | 23 | 0.496 | 1.522 | 1.026 |  | 206.9% |  |
|  | 30 | 0.562 | 9.868 | 9.306 |  | 1655.9% |  |
|  | 33 | 0.890 | 27.691 | 26.801 |  | 3011.3% |  |
|  | 34 | 0.010 | 24.083 | 24.073 |  | 240730.0% |  |
| 5M BBN<br>1M RosiTram | 7 | 4.467 | 2.190 | -2.277 | -3.198 | -51.0% | -91.0% |
|  | 9 | 11.943 | 0.315 | -11.628 |  | -97.4% |  |
|  | 12 | 2.604 | 0.000 | -2.604 |  | -100.0% |  |
|  | 15 | 5.408 | 0.000 | -5.408 |  | -100.0% |  |
|  | 20 | 0.516 | 0.000 | -0.516 |  | -100.0% |  |
|  | 21 | 0.562 | 0.000 | -0.562 |  | -100.0% |  |
|  | 25 | 0.523 | 0.000 | -0.523 |  | -100.0% |  |
|  | 13 | 1.461 | 0.000 | -1.461 |  | -100.0% |  |
|  | 16 | 10.014 | 1.241 | -8.773 |  | -87.6% |  |
|  | 14 | 6.261 | 0.000 | -6.261 |  | -100.0% |  |
|  | 24 | 1.518 | 0.507 | -1.011 |  | -66.6% |  |
|  | 8 | 1.148 | 0.329 | -0.819 |  | -71.3% |  |
|  | 18 | 0.406 | 0.000 | -0.406 |  | -100.0% |  |
|  | 28 | 2.523 | 0.000 | -2.523 |  | -100.0% |  |
| 5M BBN<br>1M Rosi only | 41 | 2.379 | 2.903 | 0.524 | 3.412 | 22.0% | 115.2% |
|  | 42 | 6.140 | 20.129 | 13.989 |  | 227.8% |  |
|  | 44 | 3.906 | 0.000 | -3.906 |  | -100.0% |  |
|  | 45 | 7.947 | 24.451 | 16.504 |  | 207.7% |  |
|  | 46 | 20.929 | 0.887 | -20.042 |  | -95.8% |  |
|  | 47 | 3.120 | 16.525 | 13.405 |  | 429.6% |  |
| 5M BBN<br>1M Tram only | 10 | 1.554 | 12.217 | 10.663 | 3.783 | 686.2% | 235.5% |
|  | 33 | 2.226 | 20.224 | 17.998 |  | 808.5% |  |
|  | 35 | 6.152 | 4.197 | -1.955 |  | -31.8% |  |
|  | 36 | 4.149 | 5.564 | 1.415 |  | 34.1% |  |
|  | 38 | 6.545 | 2.424 | -4.121 |  | -63.0% |  |
|  | 39 | 6.133 | 4.829 | -1.304 |  | -21.3% |  |

MEK/ERK signaling is an important driver of tumor survival, growth and proliferation in numerous cancers, down-stream of activating mutations in *EGFR*, *ERBB2*, *FGFR3*, *K-RAS* and BRAF ^25–27^. *Pparg* transcriptional activity can be suppressed by MEK/ERK signaling ^28–30^, and studies using mesenchymal breast cancer models showed that Trametinib, a MEK inhibitor, augmented Rosi activity, inducing tumor cells to undergo adipogenesis ^18^. We therefore tested whether Trametinib (hereafter referred to as Tram), could increase the anti-tumor activity of Rosi in BBN-induced BASQ tumors. Mice treated with BBN for 5 months were dosed either with Rosi or Tram alone or with a combination of Rosi+Tram by oral gavage for one month, and tumor growth was assessed by ultrasound imaging. In mice treated with Tram alone, we observed a small reduction in the rate of tumor growth, similar to what was observed with Rosi alone (Fig. 1C,D,E,F,I). However, in mice treated with both Rosi and Tram, tumors were decreased in size by an average of 91%; and in most cases (9/14), residual tumors were not detectable (Table 1; Fig. 1G,H,I). Histopathological evaluation confirmed the observations from ultrasound of Rosi/Tram treated tumors. Bladders from BBN-treated animals dosed with vehicle for 1 month contained large tumors, many of which were muscle invasive and exhibited signs of squamous differentiation [T2-T3; Table 2, Fig. 1J,O]. Tumors from BBN treated animals that received either Rosi or Tram alone displayed less severe histopathological changes [Table 1; (Dysplasia-T1)] compared to vehicle control tumors and lacked overt signs of squamous differentiation (Fig. 1I,K,L,P,Q). Consistent with the ultrasound results, bladders from animals treated with both Rosi and Tram contained few tumors, and the histology of the urothelium was quite similar to that in untreated healthy animals (Table 2; Fig. 1 I,M,N,R,S,). These findings together, suggest that Rosi and Tram alone have significant effects on tumor growth, but the combination of both drugs is more powerful, inducing tumor regression after one month of continuous treatment.

**Table 2.** Histopathological tumor grading based on H&E.

| Treatment | Number | Evaluation | Tumor Grading |
| --- | --- | --- | --- |
| 5M BBN<br>1M Vehicle | 3 | dysplasia with squamous keratinization |  |
|  | 5 | muscle invasive carcinoma with squamous+glandular, surface with squamous metaplasia | T3 |
|  | 13 | muscle invasive urothelial carcinoma with extensive squamous +focal glandular differentiation | T3 |
|  | 15 | invasive carcinoma with extensive squamous+ CIS | T3 |
|  | 17 | invasive carcinoma with poorly differentiated+ sarcomatous differentiation and CIS | T1 |
|  | 21 | early invasive carcinoma and CIS | T1 |
|  | 38 | invasive carcinoma+ noninvasive with extensive squamous differentiation | T1 |
|  | 76 | superficial invasive carcinoma and dysplasia+ squamous metaplasia | T1 |
| 5M BBN<br>1M RosiTram | 1 | early invasive carcinoma and CIS | T1 |
|  | 5 | normal |  |
|  | 6 | normal/ reactive atypia |  |
|  | 7 | normal |  |
|  | 8 | normal |  |
|  | 9 | atypia |  |
|  | 10 | atypia |  |
|  | 12 | atypia |  |
|  | 13 | atypia |  |
|  | 15 | normal/focal hyperplasia |  |
|  | 20 | normal/focal hyperplasia |  |
|  | 21 | normal/focal hyperplasia |  |
|  | 25 | normal/focal hyperplasia, granulomatous infiltration? |  |
| 5M BBN<br>1M Rosi | 41 | dysplasia |  |
|  | 42 | invasive carcinoma possible sarcomatous differentiation/ surface with CIS |  |
|  | 44 | atypia |  |
|  | 45 | dysplasia |  |
|  | 46 | dysplasia |  |
|  | 47 | CIS with squamous changes, extensive inflammation |  |
| 5M BBN<br>1M Tram | 10 | dysplasia |  |
|  | 33 | dysplasia |  |
|  | 35 | dysplasia+ possible granuloma |  |
|  | 36 | dysplasia |  |
|  | 38 | dysplasia |  |
|  | 39 | dysplasia+ possible granuloma |  |

As is the case in human BASQ tumors, BBN-induced tumors in mice express squamous markers including *K14* and *K6a* and lack intermediate cells and superficial cells, populations that normally express luminal markers including *Kr*t20, *Pparg*, *Foxa1* and *Gata3* (Fig. 2A,B,E-H). Analysis of the urothelium in animals treated either with Rosi or Tram alone revealed that upper levels of the urothelium, which were initially populated by BASQ cells, were populated with intermediate and superficial cells expressing appropriate luminal markers, including *Upk2* and *Krt2*0, but the basal-most layers still contained squamous cells including K6a+ K14+ cells, indicating that Rosi and Tram alone were not sufficient to respecify the basal population (Fig. 2E,I-N). In animals treated with combined Rosi and Tram, however, few residual tumors remained, and the squamous differentiation was reversed, restoring the urothelium a healthy state that was populated with normal endogenous cell types (K5-Basal cells, K14-Basal cells, intermediate cells and superficial cells (Fig. 2C,D,E,O-T).

**Figure 2.**
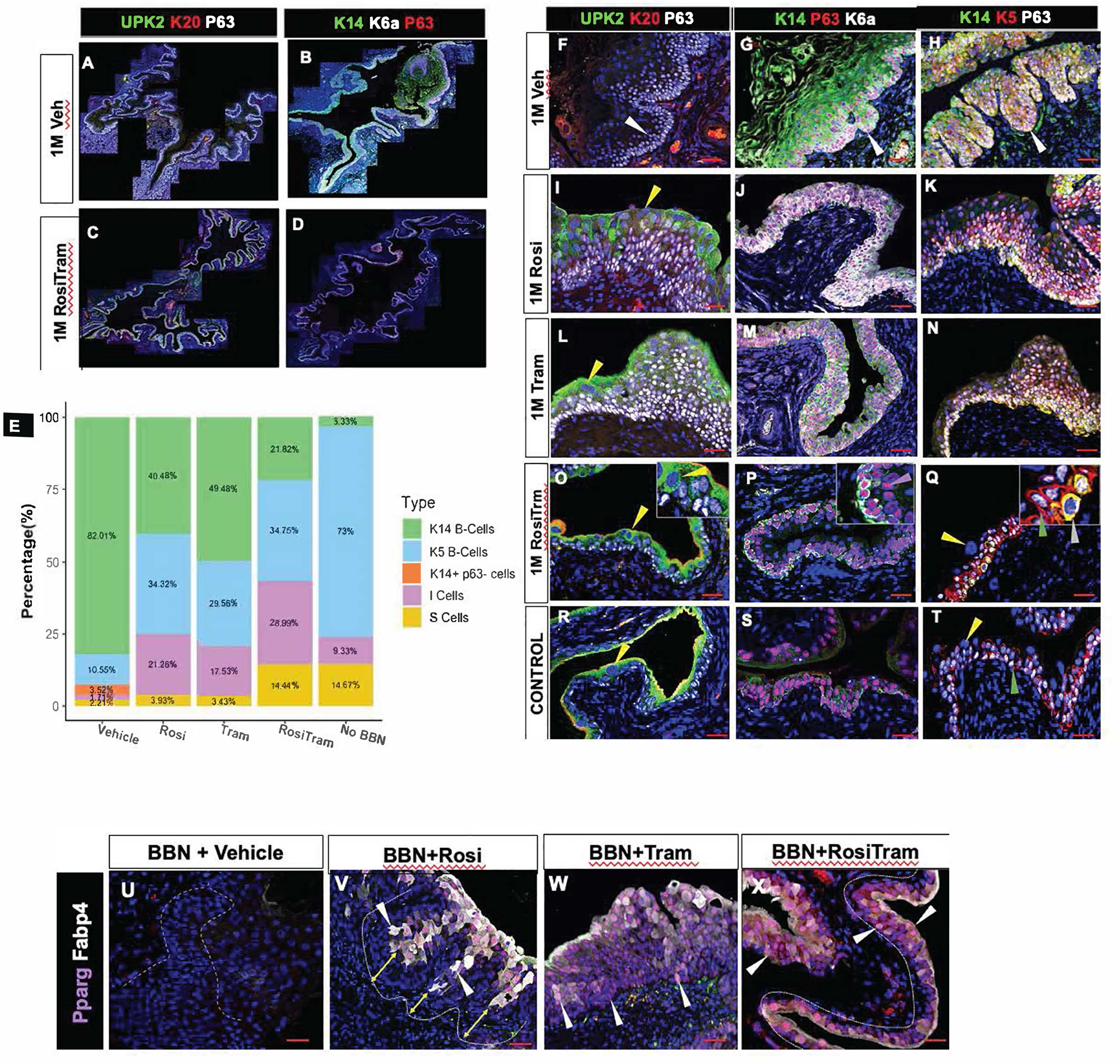
Combined Rosi+Tram treatment impairs tumor growth and restores normal urothelial differentiation in mice with BBN-induced Ba/Sq tumors. **A-D.** Panoramas showing luminal and squamous/basal marker expression in mouse exposed to BBN for 5 months then treated with vehicle **(A,B)** or Rosi+Tram **(C,D). E.** Composition of the cell types in the urothelium in animals treated with Vehicle, Rosi, Tram Rosi+Tram and no BBN controls**. F,I,L,O,R.** Upk2, K20 and P63 expression in animals treated with BBN for 5 months then treated with vehicle **(F)**, Rosi **(I)**, Tram **(L)**, Rosi+Tram **(O)** and untreated adult mouse controls **(R). G,J,M,P,S.** Expression of K14, P63 and K6a, in the urothelium of mice treated with BBN for 5 months then with Vehicle for one month (**G**), Rosi for one month (**J**), Tram for 1 month (**M**) combined Rosi/Tram for one month. (**P**) and untreated controls **(S)**. **H,K,N,Q,T**. Expression of K14, K5 and P63 in the urothelium of mice treated with BBN for 5 months then with vehicle for one month (**H**), Rosi for one month (**K**), Tram for one month (**N**), Combined Rosi/Tram for one month (**Q**), and untreated controls (**T**). U-Y. *Pparg*/Fabp4 expression in a wild type adult mouse control **(U)**, after treatment for one month with vehicle, (**V**), with Rosi (**W**), with Tram, (**X**) with Rosi+Tram **(Y**). White arrowheads in (F-H) depict the invading front of tumors; Yellow arrowheads point to superficial cells. Arrowheads in **(V-X)** indicate the boundary of *Pparg* activation in tumors and in the of treated and untreated mice, based on expression of Fabp4. Scale bars: 50μm.

To begin to examine the specific effects of Rosi and Tram, we determined their effects on the distribution of cells with active *Pparg* signaling, using expression of *Fabp4*, a direct *Pparg* transcriptional target, as a proxy. As expected, neither *Pparg* nor *Fabp4* were detectable in BASQ tumors treated with vehicle, while in tumors treated with Rosi alone, *Pparg* expression and signaling were co-localized in upper layers of invasive tumors but were absent from basal-most lower layers (Fig. 2U,V; yellow arrows denote basal-most cell layers of the tumor). Treatment with Tram alone or Tram in combination with Rosi, increased the domain of *Pparg* signaling so that it included both the suprabasal and basal-most layers (Fig. 2W,X). These observations suggest that cells in the basal-most layers of BASQ tumors, which includes invasive tumor cells and progenitors, are resistant to Rosi, but can be sensitized by MEK inhibition.

*Pparg* expression and signaling can be down-regulated by MEK/ERK dependent phosphorylation ^28–30^, and the pathway is frequently activated down-stream of Egfr, Ras, or Raf mutations in BASQ tumors in both humans and mice (Fig. S2A-D). In addition to driving proliferation and survival, MEK/ERK-dependent phosphorylation can induce SUMOylation or ubiquitination of *Pparg*, which is followed by proteasomal degradation ^31^. Recent studies identified a gene signature associated with dephosphorylation of serine 273 (S273), a MEK/ERK target site in the *Pparg* ligand binding domain that inhibits *Pparg* activation by Rosi and other *Pparg* agonists ^32,33^. RNA-seq analysis of BBN-induced tumors treated with Rosi+Tram revealed alterations in most genes in the S273 dephosphorylation signature gene set both at 4 days and 1 month after treatment, including *Aplp2*, *Car3*, *Nr1d1*, *Nr1d2*, *Nr3cs*, *Peg10*, and *Selenbp*1 (Fig. S2E). Importantly, *Gdf3*, a *Bmp* inhibitor whose expression is associated with *Pparg* dephosphorylation, was downregulated in Rosi/Tram treated tumors [Fig. 2SE; ^32^]. Together these observations indicate that the combination of Rosi and Tram, both eradicates Ba/Sq tumors and restores proper differentiation in the urothelium of mice harboring Ba/Sq tumors, which has undergone squamous metaplasia. Importantly, our studies suggest that cells residing in the basal-most layers of BASQ tumors are resistant to Rosi and can be sensitized to Rosi by addition of Tram, most likely because it reverses MEK-dependent post-translational modification of *Pparg*.

### Rosi and Tram treatment drives apoptosis and cell cycle exit in BBN-induced tumors by down-regulating the Bcl2 pro-survival pathway

The Mek-Erk-Mapk cascade drives expression of pro-survival factors including *Bcl-2*, family members in a number of tumor types ^34–39^. *Pparg* synthetic agonists have a similar effect as Tram, inducing apoptosis and down-regulating *Ccnd1* ^40–42^. To examine the mechanism by which BBN-induced tumors were suppressed following Rosi and Tram treatment, we analyzed them after 1, 4, and 7 days after administration of the 2 drugs for expression of activated Caspase-3, to identify cells undergoing apoptosis ^43^. Large numbers of apoptotic tumor cells expressing activated Caspase-3 were first observed at 7 days post-treatment; (Fig. 3A-E). In addition, we observed a marked decrease in the numbers of Ki67-positive cells in treated tumors suggesting that the two-drug combination also reduced proliferation (Fig. 3F).

**Figure 3.**
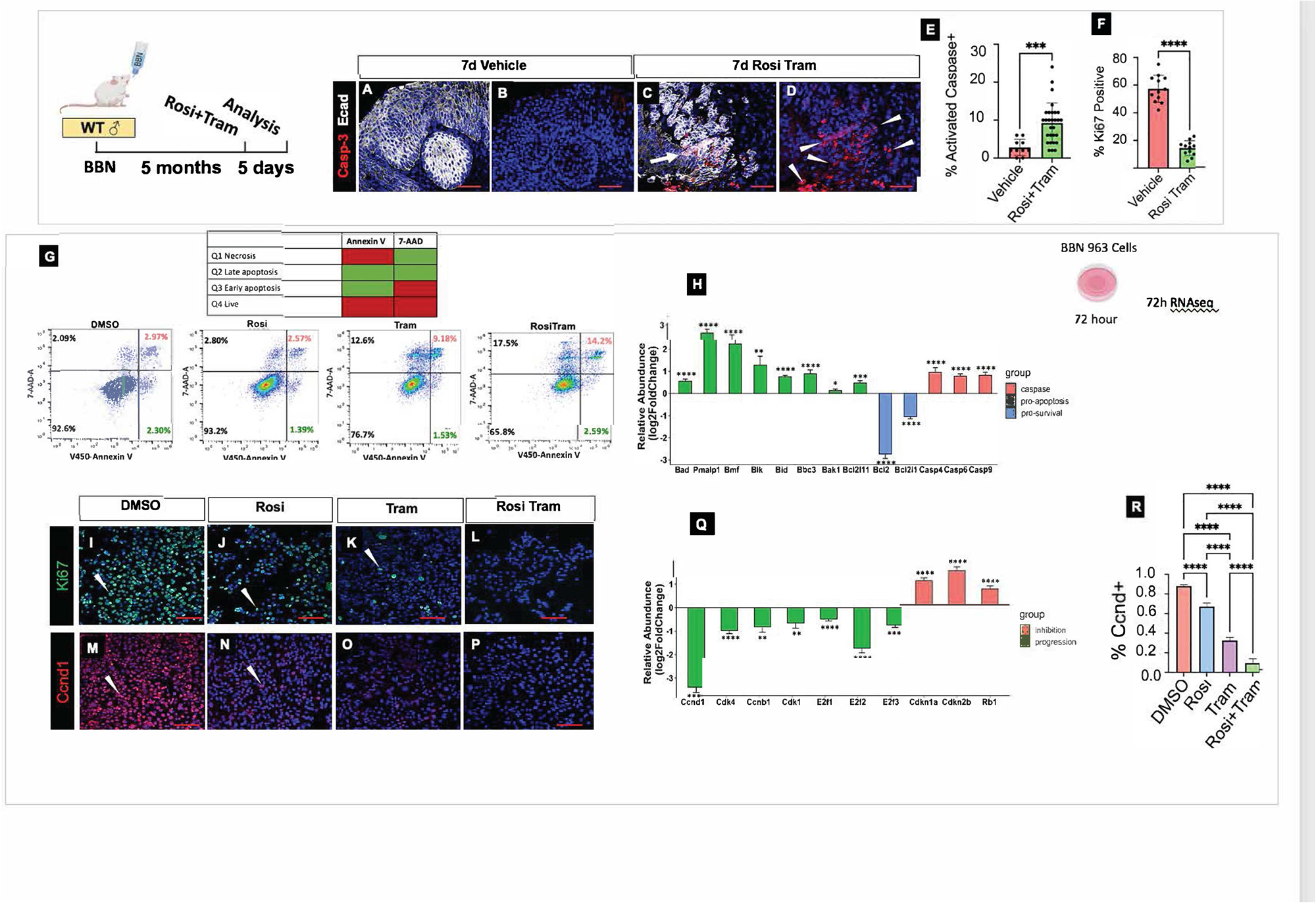
Rosi/Tram treatment induces apoptosis and reduces proliferation in BBN-induced Ba/Sq tumors after 7 days. **A,B**. Expression of activated Caspase 3 in tumors of mice treated with BBN for 5 months then with Vehicle for 7 days. **C,D.** Expression of activated Caspase 3 in tumors of mice treated with BBN for 5 months then with combined Rosi+Tram for 7 days **E**. Bar graph showing the percentage of cells expressing activated Caspase 3 in tumors from BBN-induced mice treated with vehicle for 7 days or with Rosi+Tram for 7 days. **F.** Bar graph showing the percentage of cells expressing Ki67 in animals treated with BBN for 5 months with vehicle or Rosi+Tram. **G**. Apoptosis assayed by AnnexinV-eFluor450/7-AAD double staining in BBN963 cells treated with DMSO, Rosi, Tram or combined Rosi+Tram for 72h. Cells undergoing early apoptosis were AnnexinV+/7-AAD-. Cells undergoing late apoptosis were AnnexinV+/7-AAD+. * p ≤ 0.05; ** p ≤ 0.01; *** p ≤ 0.001; **** p ≤ 0.0001. **H.** Up-regulation of proapoptotic genes and down-regulation of pro-survival genes in Rosi/Tram treated Ba/Sq cells compared to controls. **I-L.** Rosi, Decreased proliferation in BBN 963 Ba/Sq cells treated with Rosi, Tram, or combined Rosi+Tram. **M-P.** Down-regulation of Ccnd1 in BBN 963 cells after treatment with Rosi, Tram or Rosi+Tram. **Q**. Down-regulation of Ccnd1 and genes that positively regulate cell cycle progression, and up-regulation of genes that inhibit progression in BBN 963 cells treated with Rosi+Tram. **R.** Bar graph showing levels of Ccnd1 after treatment with vehicle, Rosi, Tram and combined Rosi+Tram. Scale bars: 50μm

To further investigate the individual and combined roles of Rosi and Tram, we performed in vitro and in vivo studies using BBN 963 cells, a murine bladder cancer line derived from a BBN-induced BASQ tumor ^44^. BBN 963-derived orthotopic tumors and cultured cells expressed BASQ markers including Krt6a, Krt14 (Fig. S31, A-H, I-K). Importantly, analysis of orthotopic grafts and BBN 963 cells in culture indicated that this cell line responds to Rosi and Tram in a similar way as BBN-induced Ba/Sq tumors: we observed up-regulation of luminal markers such as *Pparg* and *Upk*, and down-regulation of basal markers in both orthotopic tumor grafts and 2-D cultures of BBN 963 cells (Fig. 3S1, I-N). Annexin V staining identifies apoptotic cells that express surface-exposed phosphatidylserine (PS) and 7-AAD is a DNA intercalator that labels cells that have lost membrane integrity, a feature of late apoptosis and secondary necrosis ^45^. Analysis of BBN 963 cells treated with Rosi and Tram revealed low numbers of cells in early/late apoptosis in cultures treated for 7 days with vehicle or Rosi alone, while Tram treatment alone led to increased numbers of cells in late apoptosis (Fig. 3G). Combined treatment with Rosi and Tram resulted in further increases in the percentage of cells both in early and late apoptosis (Fig. 3G), suggesting that the 2-drug combination acts in a synergistic manner. RNA-seq analysis of Rosi+Tram treated cultures revealed significant upregulation of pro-apoptotic members of the *Bcl-2* family (*Bad*, *Noxa*, *Bmf*, *Bik*, *Bid*, *Puma*, Bak, and *Bim*), and downregulation of pro-survival members (*Bcl-2* and *Bcl-X*), known targets of both *Pparg* and MEK/ERK inhibition (Fig. 3H).

Consistent with reduced proliferation, Rosi and Tram had profound effects on the cell cycle, (Fig. 3I-R, Fig. 3S2A-H). RNA-seq analysis of BBN 963 cells after combined Rosi and Tram treatment revealed decreased expression of *Ccnd1* and other positive regulators of the cell cycle including *Cdk1*, *Cdk4*, *E2f2*, and up-regulation of negative regulators of the cell cycle, including *Rb*, *Cdkn1a* and *Cdkn2b*, both in tumors and in BBN 963 cells (Fig. 3Q; 2S3, A-H). These observations together suggest that combined Rosi+Tram treatment efficiently induced apoptosis and cell cycle arrest in BASQ tumors in vivo and in vitro, within one week of treatment.

### RA, AP-1, and Kdm6a are regulated by Rosi/Tram treatment

Paired ATAC-seq/RNA-seq analysis performed 5 days after treatment of BBN-induced tumors with either vehicle or Rosi+Tram revealed up-regulation *Pparg*-dependent pathways evidenced by increased expression of Fabp4, Plin4, Cpt2 and Cpt1a, genes important for lipid metabolism and beta oxidation that are directly regulated by *Pparg* (Fig. 4A; Fig. S4A-E). Consistent with these findings, we observed up-regulation of Cpt1a expression in the urothelium of Rosi+Tram treated mice, as well as accumulation of neutral lipid droplets, which were not detected in controls (Fig. S4A-D); alterations observed previously in mice expressing a constitutively active form of *Pparg* in the urothelium ^12^. Up-regulated pathways both in RNA-seq and ATAC-seq analysis include luminal markers (*Fgf3*, *Upk1b*, *Grhl3*, *Foxa1*, *Erbb2*) and interestingly, we also observed up-regulation of genes that control the RA-signaling pathway (Fig. 4A-D).

**Figure 4.**
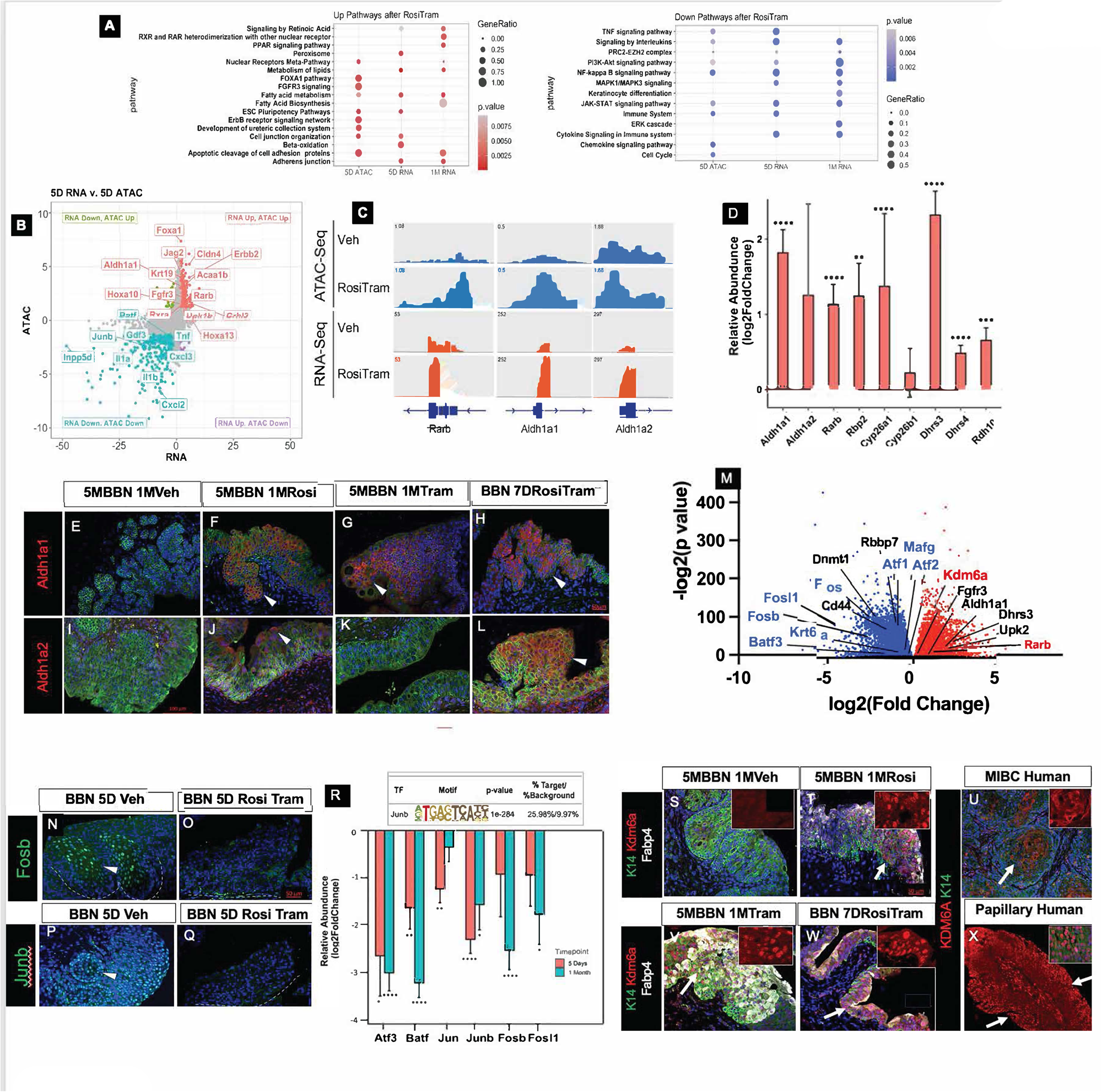
Rosi+Tram treatment induces retinoid signaling and expression of Kdm6a and decreases AP-1 in BBN-induced Ba/sq tumors. **A.** Pathways changed from ATAC-seq/RNA-seq analysis of mice treated with BBN for 5 months, then dosed with Rosi+Tram for 5 days or 1 month, compared to controls. Bubble color and size correspond to the p-value (p<0.05) and number of genes enriched in the pathway. **B**. Volcano plot showing results of combined ATAC-seq/RNA-seq analysis of tumors from mice treated with BBN for 5 months, then with Rosi+Tram for 5 days. Red indicates genes that are both more accessible and up-regulated after 5 days of RosiTram treatment; turquoise designates genes that are both less accessible and down-regulated after Rosi Tram treatment. Intersection of RNA-seq and ATAC-seq plotted as -log10(p-value) of Log2 fold change in animals dosed with Rosi+Tram compared to vehicle. Each point represents one gene. **C**. IGV tracks showing expression and chromatin accessibility in genes important for retinoid signaling in tumors from mice treated with BBN for 5 months, then with Rosi+Tram for 5 days. **D.** Bar graph showing up-regulated *Pparg* targets after Rosi+Tram treatment, based on bulk RNA-seq analysis. **E-H**. Expression of the RA-synthesizing enzyme Aldh1a1 in mice treated with BBN for 5 months, then treated with vehicle (**E**), Rosi (**F**), Tram (**G**), and combined Rosi+Tram (**H**). I-L. Expression of the RA-synthesizing enzyme Aldh1a2 in tumors from animals treated with BBN for 5 months, then dosed with Vehicle **(I)** Rosi (**J**), Tram (**K**) and Rosi+Tram (**L**). **M.** Volcano plot showing genes whose expression increased (red) and decreased (blue) following Rosi/Tram treatment of Ba/Sq BBN 963 cells. Colored points have a p-value ≤ 0.05. **N-Q.** Tumors from mice treated with BBN for 5 months then with vehicle or with Rosi, Tram or Rosi+Tram for 5 days, then stained for expression of Fosb (**N,O**) or JunB (**P,Q**).**R.** HOMER motif analysis after ATAC-seq showing enrichment of targets within the Junb regulatory network localized within less accessible peaks in animals treated with Rosi+Tram for 5 days compared to vehicle. **S-X**. Immunostained sections from mice treated with BBN for 5 months, then treated for one month with Vehicle (**S**), Rosi (**T**), Tram (**V**), Rosi+Tram (**W**) stained for expression of Krt14 (green), Kdm6a (red) and Fabp4 (white). **U**. A Ba/Sq tumor from a patient with MIBC stained for expression of Kdm6a (red). **X**. A papillary tumor from a patient stained with Kdm6a (red). Scale bars: 50μm

The retinoid signaling pathway has long been known to suppress squamous differentiation in the urothelium and other epithelia ^46–48^. Studies in mouse mutants and in human cells indicate that RA-signaling promotes urothelial/luminal differentiation, both during development and in the adult during injury induced regeneration ^49–51^. RA-regulated genes that were both increased and more accessible after Rosi/Tram treatment including *Rarb*, a canonical RA-target that harbors an RA-response element in its promoter ^52^, *Cyp26a1*, *Cyp26b1*, *Dhrs3*, and *Dhrs4*, that are also known RA-targets. Importantly, we observed up-regulation of Rdh10, a retinol dehydrogenase that converts retinol to retinaldehyde, and *Aldh1a1* and *Aldh2a*2, retinaldehyde dehydrogenases that convert retinaldehyde to retinoic acid, the active metabolite (Fig. 4B-D,M). Analysis of BBN tumors treated with Rosi, Tram or both drugs, revealed little expression of Aldh1a1 in untreated tumors, whereas expression was increased after treatment with either Tram or Rosi alone, and further increased after treatment with both drugs ((Fig. 4E-H). *Aldh1a2,* which like *Aldh1a1* is barely detectable in BASQ tumors, was increased slightly with Rosi alone and was barely detectable in Tram treated tumors but was strongly up-regulated after treatment with Rosi+Tram (Fig. 4J-L).

RNA-seq analysis of Rosi/Tram treated BBN 963 cells produced similar results. We observed up-regulation of luminal markers including *Fgfr3*, and *Upk2*, consistent with the shift from BASQ to luminal differentiation, as well as increased expression of genes regulated by RA-signaling (*Rarb, Dhrs3*) (Fig. 4M). We also observed up-regulation of *Kdm6a*, an H3K27me3 lysine demethylase that positively regulates luminal differentiation in mouse urothelial organoids ^19^. Analysis of expression of *Kdm6a* after Rosi, Tram or Rosi/Tram treatment of BBN-induced tumors revealed little or no detectable Kdm6a in untreated tumors, while expression increased with Rosi or Tram alone (Fig. 4S,T,V), and expanded to include most cells in tumors and in the urothelium after treatment with both drugs (Fig. 4W). Analysis of the distribution of *KDM6A* in BASQ and LP human tumors revealed non-nuclear expression in suprabasal layers of MIBC, while nuclear expression was widespread in LP tumors, observations that are consistent with the known role of *Kdm6*a as a promoter of luminal differentiation (Fig. 4U,X)

Pathways that were both less accessible and down-regulated after Rosi/Tram treatment of BBN-induced tumors include Tram targets (MAPK1/MAPK3 and ERK), Keratinocyte differentiation, and Nf-kb, a known *Pparg* target that is a master regulator of immunity and inflammation (Fig. 4A). HOMER motif analysis of less accessible sites after ATAC-seq analysis revealed enrichment of the *JunB* motif and down-regulation of AP-1 family members including *Jun, Batf, Atf3, Fosb* and *Fosl1* (Fig. 4R). Consistent with this, analysis of expression of Junb and Fosb in BBN-induced tumors revealed high expression of both genes in tumors treated with vehicle, while both genes were undetectable after treatment with Rosi/Tram (Fig. N-Q). Similar findings were obtained from RNA-seq analysis of Rosi/Tram treated BBN 963 cells (Fig. 4M).

*Pparg* is known to block immune infiltration in bladder cancer, at least in part via repression of *Rela*, a critical component of the Nf-kb complex ^12,53–56^. Consistent with this, we observed decreased expression of *Rela* and other components of the Nf-kb signaling pathway after Rosi/Tram treatment (Fig. 4A), and HOMER motif analysis of less accessible sites after ATAC-seq analysis revealed enrichment of *Rela* motifs within the Nf-kb network, including *Nfkb1* and *Nfkb2* (Fig. S4G), which are positively regulated by *Rela* ^57^. We observed down-regulation of interleukins, cytokines, MHC Class II and genes important for T-cell signaling following Rosi/Tram treatment (Fig. 4F). Analysis of *Rela* and CD45 is confirmed these observations. We found significant expression of Rela, and significant immune infiltration in untreated tumors. Both were relatively unaffected by treatment with Tram alone (Fig. S4H,I) and were slightly reduced after Rosi treatment (Fig. S4J,K), but they were nearly undetectable after Rosi+Tram administration (Fig. S4L-O),.

### Retinoid signaling controls Basal vs luminal differentiation down-stream from *Pparg*

Deficiency in vitamin A, the retinoid precursor, induces squamous metaplasia of epithelia lining the airways, prostate, vagina and bladder ^48,58,59^. Previous studies from our laboratory indicated that retinoid signaling was required both for formation of the urothelium in mice and for regeneration in response to injury ^60^. To investigate directly how RA-regulates growth and differentiation in BASQ tumors, we engrafted BBN 963 cells into syngeneic mice. After 2 weeks animals were dosed with RA or vehicle (DMSO) via oral gavage and assessed for the effects on tumor growth and differentiation. This analysis revealed decreased proliferation in tumors treated with RA compared to controls (Fig. 5A-D,G) and up-regulation of luminal markers (Fig. 5A). RNA-seq analysis confirmed these observations and revealed a number of shared targets regulated both by RA and Rosi/Tram, including RA-regulated genes (*Rarb*, *Cyp26b1*, *Dhrs3*, *Rbp4*, *Stra6*) luminal markers (*Fgfr3, Krt19*, *K20* and *Upks*) and *Kdm6a* (Fig. 5F,E). Down-regulated genes after both RA and Rosi/Tram treatment included BASQ markers (*Krt14, Krt16, Krt6a, CD44* (Fig 4E,F). Interestingly, both RA and RosiTram treatment resulted in decreased expression of AP-1 family of members, (*Fosb, Fosl1, Fosl2*), which are drivers of squamous differentiation and are known to oppose RA-signaling in a number of contexts, including mouse urothelial cells and organoids ^19^.

**Figure 5.**
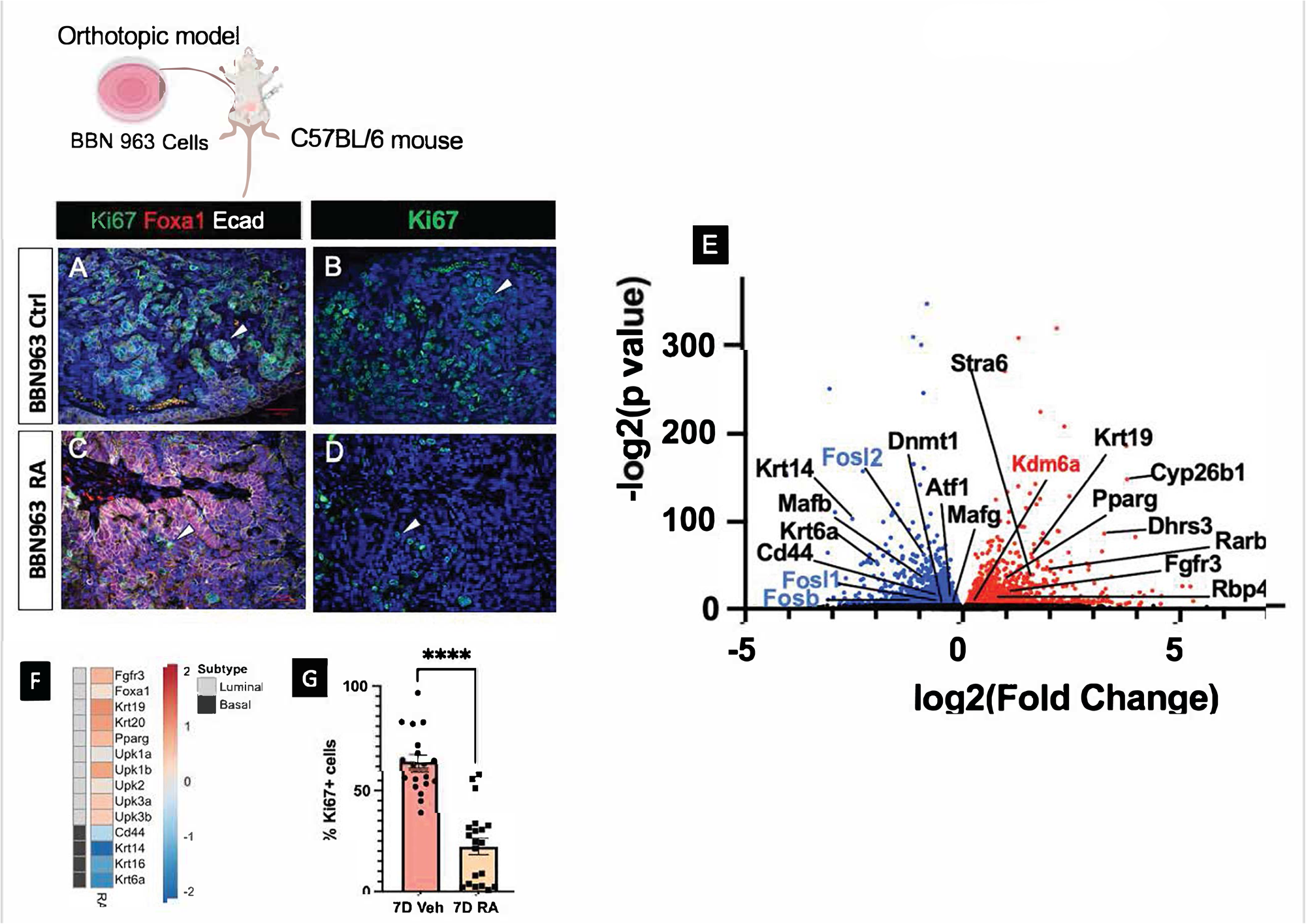
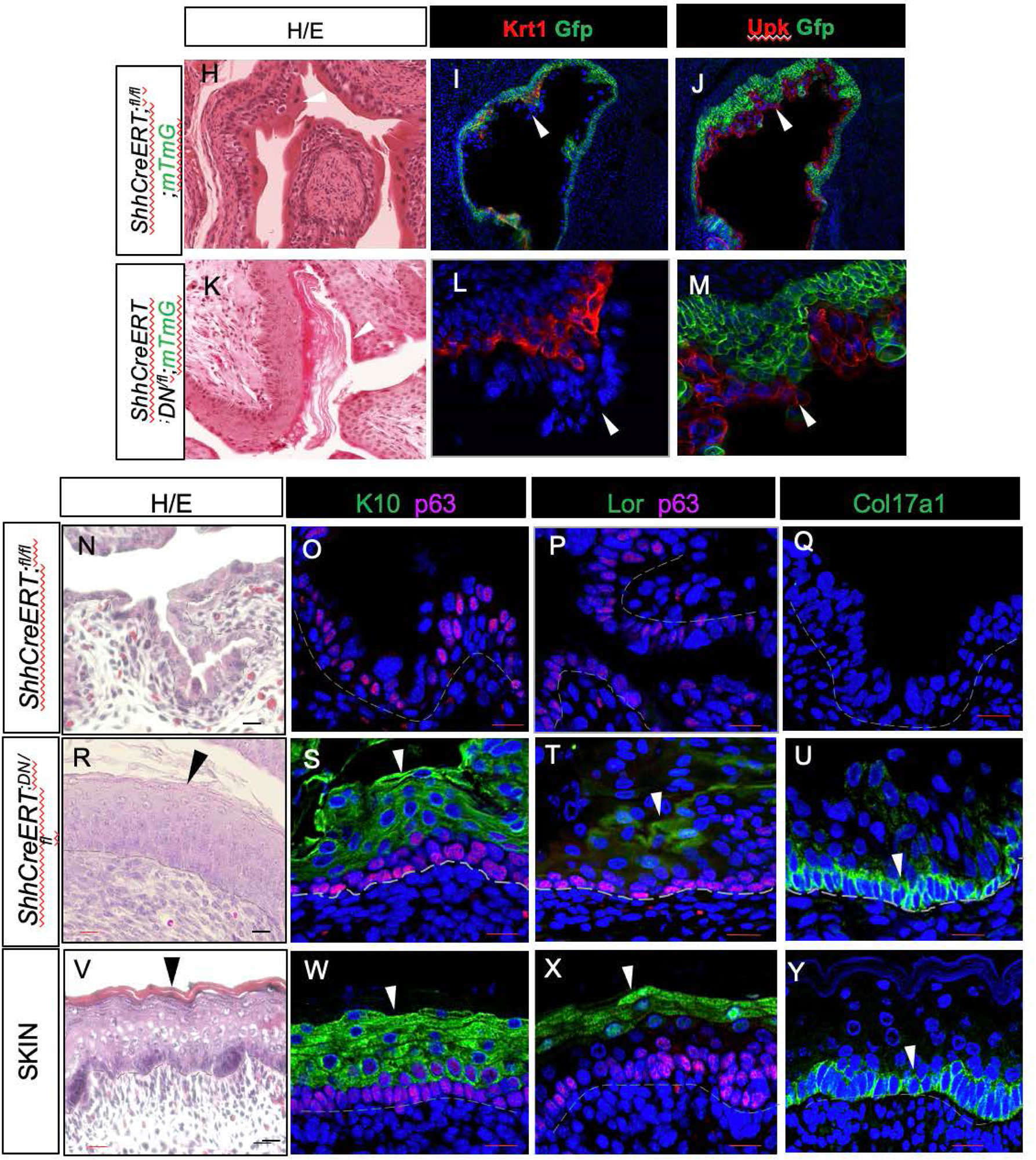
Retinoic acid induces BBN 963 cells to undergo a Basal-to-luminal shift in differentiation and lowers proliferation. **A,B.** Ki67 (green) and Foxa1 (red) staining in an orthotopic BBN 963 tumor treated with vehicle for 7 days. **C,D.** Ki67 (green) and Foxa1 (red) staining in an orthotopic BBN 963 tumor treated with RA for 7 days. **E.** Volcano plot showing changes in luminal and Ba/Sq markers, RA-regulated genes and AP-1 family members after RNA-seq analysis of BBN 963 orthotopic grafts treated wither with vehicle or RA. **F.** Changes in Ba/Sq and luminal markers after RA treatment of orthotopic BBN 963 grafts. **G.** Decreased proliferation in proliferation in BBN 963 orthotopic tumors after treatment with RA versus vehicle for 7 days. **H,K** Squamous differentiation in the bladder of control ShhCreERT2;RaraDN;mTmG mice compared to controls, 3 months after Tamoxifen induction**. I**, Lineage-marked Krt1 expressing cells in ShhCreERT2;RaraDN;mTmG of newborn mice induced with Tamoxifen at E11. **J.** Gfp+ lineage cells expressing Krt1 in the urothelium of *ShhCreERT2;RaraDN;mTmG* newborn mice induced with Tamoxifen at E11. **K.** Gfp+ lineage marked cells negative for Upk+ expression the urothelium of newborn ShhCreERT2;RaraDN;mTmG mice. Lineage-marked Krt1 expressing cells in ShhCreERT2;RaraDN;mTmG of newborn mice induced with Tamoxifen at E11. **N-Q.** Analysis of newborn control mice after staining with H/E (**N**), K10 (**O**), Lor/P63 (**P**), and Col17a1 (**Q**). **R-U.** Analysis of newborn *ShhCreERT2;RaraDN* mutants after staining with H/E (**R**), K10 (**S**), Lor/P63 (**T**) Col17a1 (**U**). **V-Y**. Analysis of newborn skin from control mice by H/E staining (**V**), expression of: K10 (**W**), Lor/P63 (**X**) and Col17a1 (**Y**). Scale bars: 50μm

Kdm6a, which was up-regulated after Rosi/Tram treatment of Ba/Sq tumors was also induced after treatment with RA (Fig. 5E). These findings are in agreement with studies suggesting that Kdm6a has an important role as a co-regulator of nuclear receptor signaling, including *Rars*, *Rxrs* and *Pparg* ^61^. Similar findings have been observed in mouse and human urothelial cells ^19^. Interestingly, despite the increase in *Kdm6a*, we did not observe changes in H3K27 methylation (data not shown) consistent with the suggestion that the *Kdm6a*’s effects on luminal differentiation do not require its methylase activity.

Vitamin A deficiency has been shown to induce squamous metaplasia in the mouse urothelium, generating a skin-like epithelial sheet that lacks endogenous urothelial populations ^62^. We previously generated a mouse model of RA-deficiency by expressing a dominant inhibitory form *of Rara* (*RaraDN*) that inhibits RA-signaling globally and selectively, without affecting other nuclear receptor signaling pathways ^49,63–65^. *RaraDN*, which is inserted in the Rosa26 locus, is activatable by Tamoxifen in cells expressing Cre recombinase. To characterize the role of retinoids in the adult urothelium, we crossed *RaraDN* mutants with the Tamoxifen inducible *ShhCreERT2* driver ^66^ and the *mTmG* reporter ^67^, to label mutant and control (*ShhCreERT2;mTmG^fl/fl^*) urothelial cells and their descendants. *ShhCreERT2;RaraDN^fl/fl^;mTmG* mice display a number of severe defects outside of the bladder, and hence do not survive long after birth. We therefore either induced expression of *RaraDN* just after birth and then analyzed urothelial differentiation after 3 months, or we induced expression during development and analyzed urothelial differentiation just after birth. These experiments revealed that keratinized squamous epithelium had replaced the urothelium in adult *ShhCreERT2;RaraDN^fl/fl^* mice (Fig. 5H,K). The mutant epithelium lacked endogenous urothelial cell types and was instead populated by cells expressing squamous markers including Krt10, Loricrin, and Col17a1, which are present in skin, but are not detected in the healthy urothelium (Fig. 5N-Y).

To better characterize the process by which the urothelium became squamous, we induced expression of the *ShhCreERT2/RaraDN;mTmG* transgene during urothelial development (at E11) then analyzed animals before birth, at E17. At this stage, we observed foci of Gfp-labeled basal cells in mutants that were positive for K14 and K1, indicating that these likely contain mutant *ShhCreERT2/RaraDN;mTmG* mutant progenitor cells that are undergoing squamous differentiation (Fig. 5I,L). Importantly, based on expression of the Gfp lineage marker, the expanding squamous cell population was negative for luminal markers including *Upk* (Fig. 5J,M). This observation indicates that the population forms beneath the endogenous urothelium, most likely as a consequence of altered specification of basal progenitors, a process that has been described in other models of vitamin A deficiency ^68^. These results indicate that retinoids are likely to be important drivers of the luminal differentiation program and suppressors or squamous differentiation, down-stream of *Pparg* and MEK signaling.

## DISCUSSION

Th integration of neoadjuvant chemotherapy with radical cystectomy in the early 2000’s improved the survival of patients with MIBC, but little substantive progress has been made since that time. Immunotherapy likely has a role in the management of this disease, but the optimal integration of this modality into the care of these patients is still a work in progress, and as such there is still a pressing need to explore therapy targeting other biological vulnerabilities of MIBC Adamo et al., 2005; Als et al., 2008; Dash et al., 2008; Dogliotti et al., 2007^69^). Numerous studies reported that MIBC display a BASq-phenotype, whereas non-invasive tumors tend to be luminal/papillary. *Pparg* is a likely driver of luminal/papillary BC has been suggested to act as a tumor suppressor in Basal/Squamous tumors. Here we sought to test the efficacy of a combination of Rosiglitazone, a *Pparg* synthetic agonist and Trametinib, a MEK/ERK inhibitor in preclinical models of MIBC. We found that BBN-induced tumors under apoptosis within 7 days of treatment, resulting in a 90% reduction in the overall tumor burden after a month of treatment. In addition to killing tumors, the combined drug treatment restored normal urothelial differentiation, inducing a shift from a BASQ to a luminal differentiation program (Fig. 6). We observed up-regulation of the retinoid signaling pathway, which is known to suppress squamous differentiation, and down-regulation of AP-1, a driver of squamous differentiation. We found that *Kdm6a*, which facilitates nuclear receptor signaling and is undetectable in BASQ tumors, was activated following treatment with Rosi/Tram (Fig. 6). These studies suggest that Rosi/Tram combination may be effective in a clinical setting for treatment of MIBC.

**Figure 6.**
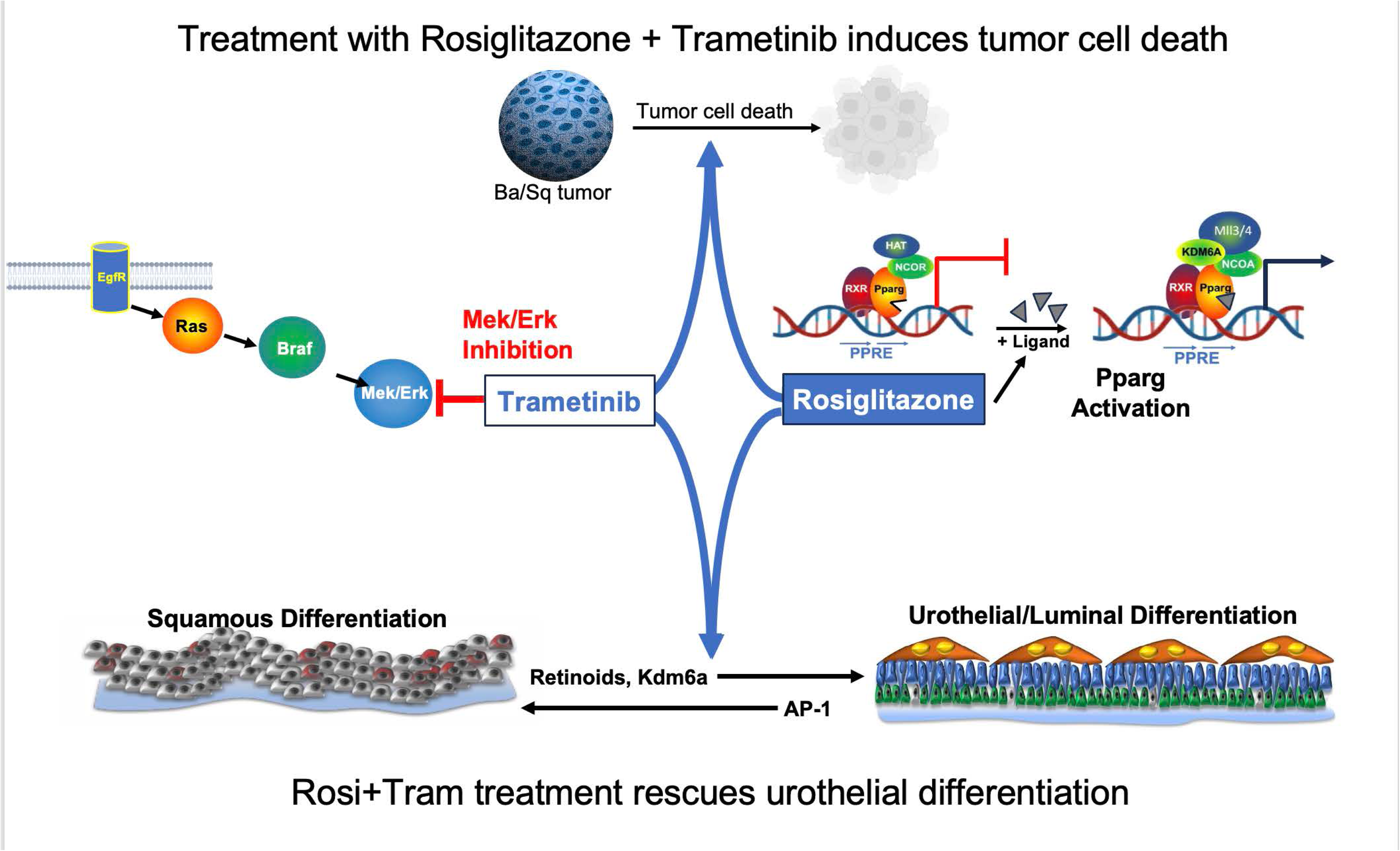
Rosiglitazone plus Trametinib treatment both eradicates tumors and restores normal urothelial differentiation in the urothelium of tumor bearing mice. Top: The respective effects of Tram and Rosi on the MEK/ERK pathway and the *Pparg* signaling pathway. Bottom: Illustration of the Ba/Sq to luminal shift in urothelial differentiation induced down-stream of Rosi+Tram, including up-regulation of retinoid signaling and Kdm6a, drivers of luminal differentiation, and down-regulation of AP-1, a driver of squamous differentiation. Bars=

### The basal-and suprabasal layers of BASQ tumors contain distinct cell populations with different drug sensitivities

*Pparg* can be activated by pharmacological inhibition of the MEK/ERK pathway ^30–33,70–73^ which is active in MIBCs downstream from mutations in *EGFR, ERBB2, FGFR3*, *K-RAS* and *BRAF* ^25–27^. Trametinib, the MEK/ERK inhibitor used in our studies has been shown to reverse MEK-dependent posttranslational modifications of *Pparg* ^74^. Consistent with this, Rosi/Tram treated tumors express a gene signature associated with MEK/ERK dependent activation of *Pparg*^33^, suggesting that Tram may activate *Pparg* a similar manner, preventing or reversing post-translational modifications. We found that Rosi treatment induced *Pparg* as well as *Fabp4*, a direct transcriptional target, in suprabasal layers of tumors populated by cells that are relatively differentiated, but not in basal-most layers of tumors, which are populated by K14-positive basal cells that are proliferative and invasive (Fig. 2U-X). That Tram alone reversed this inhibition, inducing expression of *Pparg* and *Fabp4* throughout both basal and suprabasal layers of tumors, suggests that BASQ tumors contain distinct basal and suprabasal populations, with different gene expression profiles; an observation supported by our analysis, showing differential expression of genes including *Kdm6a* and *Fosb*, both of which are expressed suprabasal but not in basal-most layers (Fig. 4).

### Basal-to-luminal differentiation in Rosi/Tram treated tumors is associated with decreased AP-1 and increased *Pparg* and RA signaling

We observed down-regulation of AP-1 family members in BASQ tumors and in cultured BASQ cells in response to Rosi/Tram treatment or retinoids (Fig. 4). The AP-1 complex is positively regulated by MAPKs and negatively regulated by nuclear receptors, including *Rars, Rxrs* and *Pparg* ^75–77^. AP-1 is known to promote Basal-squamous differentiation in the skin and in cancers ^78,79^ as well as in urothelial cells ^19^; while *Pparg* and Retinoids are known to suppress squamous differentiation in the developing urothelium, in ES cells and normal human urothelial cells ^49–51^. It will be interesting to determine whether the combination of retinoids and Tram has a similar effect on tumor survival and differentiation as observed here.

### *Pparg* and retinoid signaling promote luminal differentiation in cooperation with Kdm6a

We show here that luminal differentiation induced by either Rosi or retinoids, was accompanied by up-regulation of *Kdm6a* (*UTX*), a H3 K27 lysine demethylase that is often mutated in bladder cancer ^80^. *Kdm6a* has been shown to be necessary for RA-induced differentiation of leukemia cells ^81^, and more recently, has been shown to drive the luminal differentiation program in by promoting RA-signaling in urothelial cells ^19^. Rars are transcriptional repressors in the absence of ligand (RA) and are bound to co-repressors including N-CoR and SMRT ^82^. RA binding to Rar induces a conformational change, resulting in release of co-repressors and recruitment of co-activators including Ncoa6, which is part of the MLL3/MLL4 complex. Kdm6a interaction with Ncoa6, is thought to be important for RA-dependent transcription by recruiting the MLL3/MLL4 complex to promoters and enhancers where Rar/Rxr heterodimers are bound ^83^. Consistent with this, loss of or impaired retinoid signaling, inactivation of *Pparg* or inactivation of *Kdm6a* result in BASQ differentiation and loss of luminal urothelial populations.

### RA or *Pparg* agonists in combination with trametinib may be efficacious in treating BASQ bladder cancers

Our studies show that a combination of *Pparg* activation and MEK inhibition induced by rosiglitazone and trametinib respectively, is a powerful treatment both eradicating BASQ tumors and restoring endogenous urothelial populations, shifting the urothelial differentiation program from BASQ to luminal. Trametinib is FDA approved and is used to treat Melanoma ^74^. Rosi is also FDA approved, and has been extensively used in the past as an insulin sensitizer in patients with Type 2 Diabetes ^16^. Due to its ability to promote differentiation and cell cycle arrest, Rosi and other *Pparg* agonists have also been studied in anti-tumor therapeutics ^72,84–94^. Our studies indicate that rosi combined with trametinib is a potentially powerful treatment for BASQ bladder cancers.

## STAR Methods

### Mice

All work with mice was approved by and performed under the regulations of the Columbia University Institutional Animal Care and Use Committee. Animals were housed in the animal facility of Irving Cancer Research Center, Columbia University. Animals were housed in standard cage of 75 square inches at or below maximum cage density permitted by IACUC protocol. Temperature was maintained between 68-79°F. Humidity was maintained between 30-70%. A timed-controlled lighting system was used for a uniform diurnal lighting cycle.

### Cell lines

BBN963 cell line (W. Kim, University of North Carolina at Chapel Hill) were cultured inside tissue culture-treated plates with Dulbecco’s modified Eagle medium high glucose (DMEM) with 10% fetal bovine serum (Gibco) and 1% penicillin-streptomycin (100 U/ml; Gibco) at 37°C and 5% CO_2_. Cells were split twice per week, and cell viability was measured using trypan blue staining in the Countess II automated cell counter (Thermo Fisher Scientific).

### Orthotopic model

BBN963 cells were detached from tissue culture plates using 0.25% trypsin-EDTA at 37°C. The cells were injected orthotopically under ultrasound guidance into the bladder lamina propria of C57BL/6J male mice at 5×10^6^ cells suspended in 500uL sterile PBS using 30G needle with syringe.

### BBN Treatment

BBN (0.05%; Sigma cat#B8061-1G) was administered in the water supply *ad libidum* for up to 24 weeks to induce bladder cancer. Mice were euthanized between 20 and 24 weeks. All bladders were removed and embedded for sectioning and staining.

### Rosiglitazone and Trametinib Treatment

*In vivo.* Tumor growth was confirmed via ultrasound (Visualsonics VEVO 3100), and wild type male mice were divided into four treatment groups as follows: Group 1: Control treated with vehicle (0.5% hydroxypropyl methylcellulose and 0.2% Tween 80 in distilled water and 0.7% DMSO). Group 2: Rosiglitazone (Adipogen CAS# 122320-73-4; 20mg/kg) dissolved in vehicle. Group 3: Trametinib (LC laboratories T-8123; 0.3mg/kg) dissolved in vehicle. Group 4: Rosiglitazone (20mg/kg) and Trametinib (0.3mg/kg) dissolved in vehicle. Mice were treated daily via oral gavage for 4 days, 5 days, 7 days, or 1 month. Bladders were removed for embedding, RNAseq, or ATACseq.

*In vitro*. BBN963 cells were seeded at density of 20,000 cells/cm^2^ and treated for 24 or 72h. The cells were divided into four treatment groups as follows: Group 1: DMSO control. Group 2: Rosiglitazone (2µM). Group 3: Trametinib (100µM). Group 4: Rosiglitazone (2µM) and Trametinib (100µM). Triplicates of each treatment group were performed.

### All-trans retinoic acid Treatment

*In vitro*. BBN963 cells were seeded at density of 20,000 cells/cm^2^. DMSO control or ATRA (1*µM) (Sigma Cas# 302-79-4) dissolved in DMSO were added for 24 or 72h. Triplicates of each treatment group were performed.

### Ultrasound

Ultrasound images and scanning were acquired using VEVO 3100 Ultrasound Imaging System (FUJIFILM VisualSonics, Toronto, Canada) located within the mouse barrier in the Herbert Irving Cancer Center Small Animal Imaging facility. Tumor volume was calculated via 3D reconstruction program (Vevo LAB).

### Apoptosis Assay

The BBN963 cells were harvesting using 0.25% trypsin-EDTA at 37°C. AnnexinV/7-AAD detection was performed according to manufacturer’s instructions (eBioscience 88800672). Flow cytometry was performed on Sony MA900 Multi-Application Cell Sorter.

### RNA-Seq

For *in vivo* experiments, single cell suspensions from healthy control, vehicle control, and RosiTram treated bladders were filtered through a 70μm filter (Fisherbrand cat#22363548) and then sorted on a BD Aria II Cell Sorter using a 130 μm nozzle aperture and 13 psi pressure to collect DRAQ5+, DAPI-negative live urothelial cells. Gating strategy was performed on BD FACSDiva Software v 8.0. Cells were then centrifuged at 500 x g for 30 min at 4°C. The supernatant was discarded, and the pellet was processed for total RNA extraction. Samples with a RIN (regulation identification number) >8 were used for RNA-seq. The libraries were prepared using the SMART-Seq® v4 Ultra® Low Input RNA Kit for Sequencing (TaKaRa) followed by Nextera XT (Illumina), both according to manufacturer’s instructions. They were sequenced to a targeted depth of 40M 2×100bp reads on a NovaSeq 6000 (Illumina).

For *in vitro* experiments, total RNA was extracted and samples with a RIN >8 were used for RNA-seq. The libraries were prepared using Illumina Stranded mRNA prep (Illumina 20040532) according to manufacturer’s instruction. They were sequenced to a targeted depth of 400M 2×75bp reads on a NextSeq 550 (Illumina).

### RNA-seq Data Analysis

Differential expression analysis was performed by reading kallisto counts files into R using the R packages tximport (v.1.10.1) and biomaRt (v.2.34.2) and running DESeq2 (v.1.18) to generate log fold change values and p-values between the two experimental groups. The heatmap and PCA plots were visualized after transforming the counts using VST (variance stabilizing transformation). Gene set analysis by ConsensusPathDB (Kamburov, A. et al. 2013) were used to identify significantly changed pathways.

### ATAC-seq

Chromatin accessibility assays utilizing the bacterial Tn5 transposase were performed as described (<u>Corces et al., 2016</u>) with minor modifications. Cells (3.0 × 10^5^) were lysed and incubated with transposition reaction mix containing PBS, 1% Digitonin, Tween-20, and Transposase (llumina). Samples were incubated for 30 minutes at 37°C in a thermomixer at 1000rpm. Prior to amplification, samples were concentrated with the DNA Clean and Concentrator Kit-5 (Zymo). Samples were eluted in 20uL of elution buffer and PCR-amplified using the NEBNext 2X Master Mix (NEB) for 10 cycles and sequenced on a NextSeq 500 (Illumina).

### ATAC-seq Data Analysis

ATAC-seq reads were mapped to the mouse genome assembly mm10 using HISAT2 (v2.1.0, parameter: -X 2000). Potential PCR duplicates were removed by the function “MarkDuplicates” (parameter: REMOVE_DUPLICATES=true) of Picard (v2.23.1). The correlation analysis for genomic distribution of ATAC-seq signals was performed by the functions “multiBigwigSummary” (parameter: --binSize 1000) and “plotCorrelation” (--corMethod pearson -- skipZeros) of deepTools (v3.3.2). Peaks of ATAC-seq data were called using macs2 (v2.1.2, default parameters) and were annotated by the R package “ChIPseeker”. The distribution of ATAC-seq signals near promoter regions were visualized by the functions “computeMatrix” and “plotProfile” of deepTools (v3.3.2). The reads number for each peak was measured by featureCounts (v1.6.1). The differential accessibility of promoters was calculated by the R packages DESeq2 (v1.28.0) and visualized by ggplot2 (v3.2.1).

### Statistics and Reproducibility

All quantitation was performed on at least three independent biological samples, using the ImageJ software. Data presented in box plots are mean values ± s.e.m. Statistical analysis was performed using the R version 4.0.4. In two group comparisons, statistical significance was determined using T-test with a value of p <0.05. The number of samples used in the experiments is included in figure legends. All immunostainings and H&E experiments were performed with at least 3 biological replicates and 3 technical replicates for each condition.

### Immunostaining

Bladders were embedded in paraffin and serial sections were generated. For immunohistochemistry, paraffin sections were deparaffinized using HistoClear and rehydrated through a series of Ethanol and 1× PBS washes. Antigen retrieval was performed by boiling slides for 15 min in pH 9 buffer or 30 min in pH 6 buffer. Primary antibodies in 1% horse serum were incubated overnight at 4 °C. The next day, slides were washed with PBST three times for 10 min each and secondary antibodies were applied for 90 minutes at room temperature. DAPI (4′,6-diamidino-2-phenylindole) was either applied as part of the secondary antibodies cocktail or for 10 min, for nuclear staining, and then the slides were sealed with coverslips. Conditions of antibodies used is detailed in Table below.

### Fluorescent microscopy

Zeiss Axiovert 200M microscope with Zeiss Apotome were used to collect immunofluorescent images. Bright-field images were collected using a Nikon Eclipse TE200 microscope. Data was analyzed using the Fiji package of ImageJ (v.1.0) and Photoshop screen overlay method (v. 21.1.0).

### Bioinformatic analysis and visualization of *PPARG* and EGFR mutations in bladder cancer

This analysis was performed with R packages including “ggplot2”^95^ to generate jitter plots, “ComplexHeatmap” ^96^ and “circlize”^97^ to generate heatmap and “GSVA” ^98^ to calculate GSVA enrichment scores. Gene sets for BIOCARTA_MAPK_PATHWAY and BIOCARTA-ERK-PATHWAY were downloaded from molecular signature database in GSEA website.

### Cartoon schematics

Schematics were adapted from “Oral Tolerance Experiment”, by BioRender.com (2021). Retrieved from https://app.biorender.com/biorender-templates

| ANTIBODIES | SOURCE | IDENTIFIER | CLONE | DILUTION |
| --- | --- | --- | --- | --- |
| Chicken Polyclonal Anti-Keratin 14 | Biolegend | Cat#906001 | N/A | 1:400 |
| Chicken Polyclonal Anti-Keratin 5 | Biolegend | Cat#905901 | N/A | 1:400 |
| Rabbit Polyclonal Anti-Krt6A | LSBio | Cat#LS-B12036-100 | N/A | 1:2000 |
| Rabbit Polyclonal Anti-Keratin 10 | Covance | Cat#PRB-159P | N/A | 1:500 |
| Mouse Monoclonal Anti-Cytokeratin 20 | AgilentDako | Cat# M701929-2 | K<sub>s</sub>20.8 | 1:200 |
| Rabbit Polyclonal Anti-PPARG | Cell Signaling Technology | Cat# 2435 | N/A | 1:200 |
| Goat Polyclonal Anti-FABP4 | R&D Systems | Cat# AF1443 | N/A | 1:1000 |
| Mouse Monoclonal Anti-FOXA1 | Seven Hills Bioreagents | Cat# WMAB-2F83 | 2F83 | 1:1000 |
| Rabbit Polyclonal Anti-p63 | GeneTex | Cat#GTX102425 | N/A | 1:300 |
| Goat Polyclonal Anti-p63 | R&D Systems | Cat#AF1916 | N/A | 1:200 |
| Rat Monoclonal Anti-CD45 | BD Bioscience | Cat#550539 | 30F11 | 1:100 |
| Rabbit Polyclonal Anti-Ki67 | Abcam | Cat#ab15580 | N/A | 1:200 |
| Rabbit Monoclonal Anti-Cyclin D1 | Abcam | Cat#ab16663 | SP4 | 1:200 |
| Rabbit Polyclonal Anti-Active Caspase-3 | Promega | Cat#G7481 | N/A | 1:300 |
| Goat Polyclonal Anti-E-Cadherin | R&D Systems | Cat#AF748 | N/A | 1:400 |
| Rabbit Polyclonal NFkB p65 | Abcam | Cat#AB19870 | N/A | 1:300 |
| Alexa Fluor 488 Donkey Anti-Rabbit IgG | Jackson ImmunoResearch | Cat#711-545-152 | N/A | 1:700 |
| Alexa Fluor 488 Donkey Anti-Mouse IgG | Jackson ImmunoResearch | Cat#711-545-150 | N/A | 1:700 |
| Alexa Fluor 488 Donkey Anti-Chicken IgG | Jackson ImmunoResearch | Cat#703-545-155 | N/A | 1:700 |
| Alexa Fluor 488 Donkey Anti-Goat IgG | Jackson ImmunoResearch | Cat#705-545-003 | N/A | 1:700 |
| Cy3 Donkey Anti-Rabbit IgG | Jackson ImmunoResearch | Cat#711-165-152 | N/A | 1:700 |
| Alexa Fluor 594 Donkey Anti-Mouse IgG | Jackson ImmunoResearch | Cat#715-585-151 | N/A | 1:700 |
| Alexa Fluor 594 Donkey Anti-Chicken IgG | Jackson ImmunoResearch | Cat#703-585-155 | N/A | 1:700 |
| Alexa Fluor 594 Donkey Anti-Goat IgG | Jackson ImmunoResearch | Cat#705-585-147 | N/A | 1:700 |
| Alexa Fluor 647 Donkey Anti-Mouse IgG | Jackson ImmunoResearch | Cat#715-605-150 | N/A | 1:400 |
| Alexa Fluor 647 Donkey Anti-Rabbit IgG | Jackson ImmunoResearch | Cat#711-605-152 | N/A | 1:400 |
| Alexa Fluor 647 Donkey Anti-Chicken IgG | Jackson ImmunoResearch | Cat#703-605-155 | N/A | 1:400 |
| Alexa Fluor 647 Donkey Anti-Goat IgG | Jackson ImmunoResearch | Cat#705-605-003 | N/A | 1:400 |

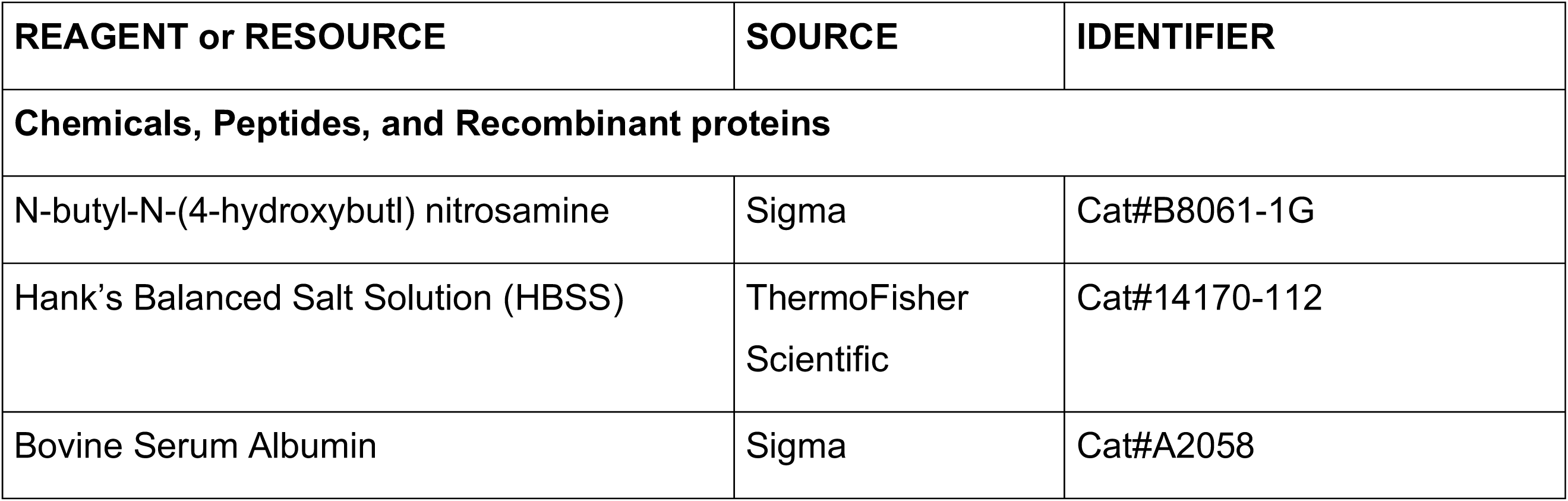
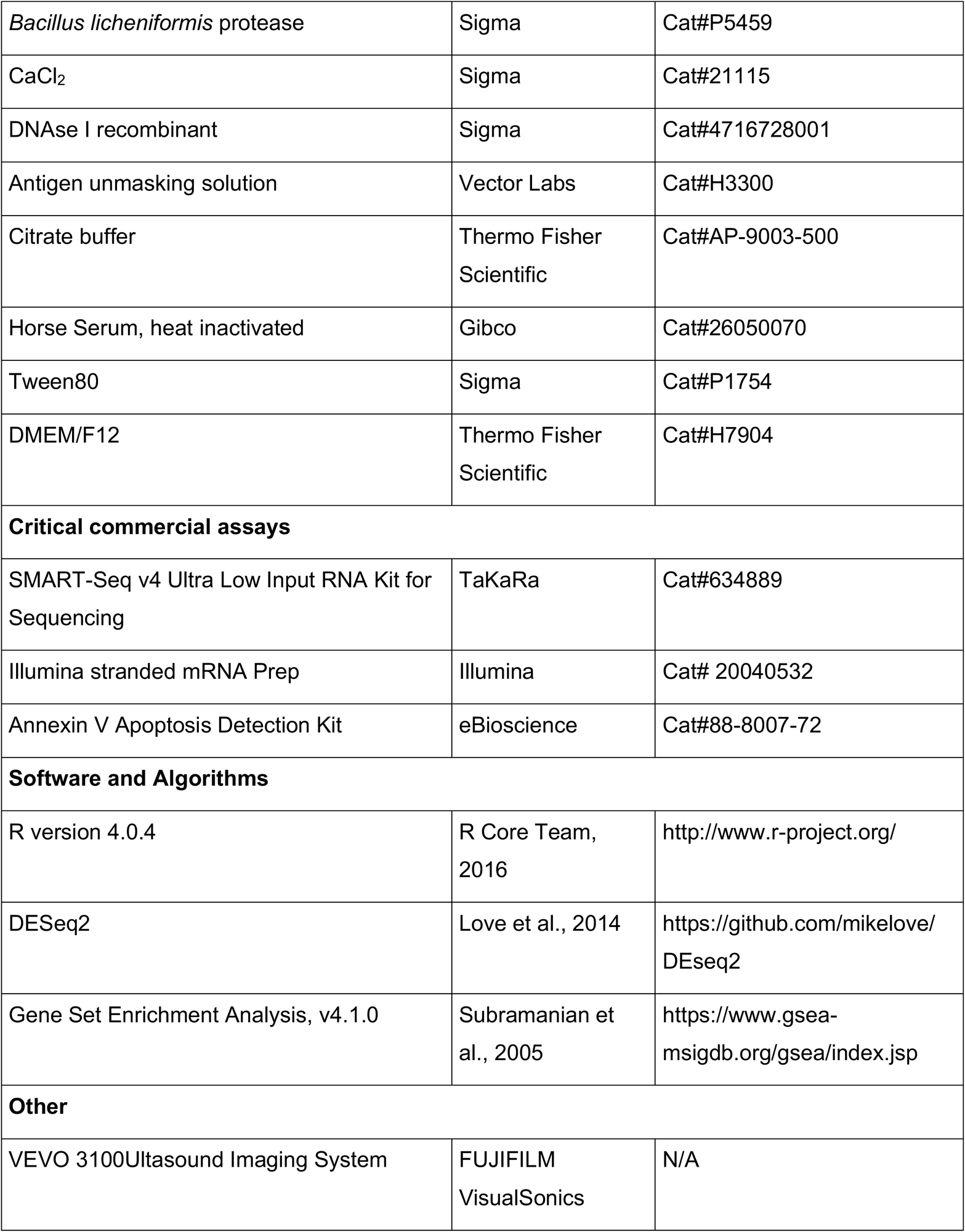

| REAGENT or RESOURCE | SOURCE | IDENTIFIER |
| --- | --- | --- |
| **Chemicals, Peptides, and Recombinant proteins** |  |  |
| N-butyl-N-(4-hydroxybutyl) nitrosamine | Sigma | Cat#B8061-1G |
| Hank's Balanced Salt Solution (HBSS) | ThermoFisher Scientific | Cat#14170-112 |
| Bovine Serum Albumin | Sigma | Cat#A2058 |
| *Bacillus licheniformis* protease | Sigma | Cat#P5459 |
| CaCl<sub>2</sub> | Sigma | Cat#21115 |
| DNAse I recombinant | Sigma | Cat#4716728001 |
| Antigen unmasking solution | Vector Labs | Cat#H3300 |
| Citrate buffer | Thermo Fisher Scientific | Cat#AP-9003-500 |
| Horse Serum, heat inactivated | Gibco | Cat#26050070 |
| Tween80 | Sigma | Cat#P1754 |
| DMEM/F12 | Thermo Fisher Scientific | Cat#H7904 |
| **Critical commercial assays** |  |  |
| SMART-Seq v4 Ultra Low Input RNA Kit for Sequencing | TaKaRa | Cat#634889 |
| Illumina stranded mRNA Prep | Illumina | Cat# 20040532 |
| Annexin V Apoptosis Detection Kit | eBioscience | Cat#88-8007-72 |
| **Software and Algorithms** |  |  |
| R version 4.0.4 | R Core Team, 2016 | http://www.r-project.org/ |
| DESeq2 | Love et al., 2014 | https://github.com/mikelove/DESeq2 |
| Gene Set Enrichment Analysis, v4.1.0 | Subramanian et al., 2005 | https://www.gsea-msigdb.org/gsea/index.jsp |
| **Other** |  |  |
| VEVO 3100Ultrasound Imaging System | FUJIFILM VisualSonics | N/A |

### Lead Contact

Further information and requests for resources and reagents should be directed to and will be fulfilled by the Lead Contact, Cathy Lee Mendelsohn.

### Data and Code Availability

Original and source data for RNAseq and ATACseq in the paper are available at GEO: https://www.ncbi.nlm.nih.gov/geo/query/acc.cgi?acc=GSE234327

## Acknowledgements

We thank William Kim for providing the BBN963 cell line, Mathieu Rouanne for data interpretation, Hyunwoo Kim and Chevaughn Waller for mouse husbandry, Yinglu Li for providing advice in ATACseq experiments. We thank Li Qiang for advice and critical reading of the manuscript. This work was supported by, U54DK1043, R01 DK095044, the JPB Foundation (C.L.M.). and T32 Training Grant DK07328 (S.A.P.). This research used the resources of the Herbert Irving Comprehensive Cancer Center Flow Cytometry Shared Resources, Molecular Pathology Shared Resources, Genomics and High Throughput Screening Shared Resources, and Oncology Precision Therapeutics and Imaging Core funded in part through Center Grant P30CA013696. Flow cytometry and cell sorting experiments were performed in the Columbia Stem Cell Initiative Flow Cytometry core facility at Columbia University Irving Medical Center under the leadership of Michael Kissner.

## Author Contributions

T.T. conceived the idea behind this study and performed the first round of experiments. T.T. also prepared RNAseq samples from the 1M RosiTram samples; S.A.P. assisted in AnnexinV/7-AAD assay and immunostaining, RNA-seq analysis of Rosi/Tram and RA-treated BBN 963 cells and tumors as well as statistical analysis. H.A. performed pathological grading of tumors; X.C., K.G., A.D.V, C.L and B.A helped with ATAC-seq data analysis and assisted in preparing sequencing figures; A.M. and G.W. performed experiments related to the *RaraDN* mutant; E.B. performed immunostaining; W,Y, helped with analysis TCGA data analysis; J.M., B.G, C.D, D.J.M. provided advice on studies and manuscript; C.L.M. helped design experiment, interpret results, and wrote the paper.

## SUPPLEMENTARY FIGURES

**Supplementary Figure Related to Figure 2.**
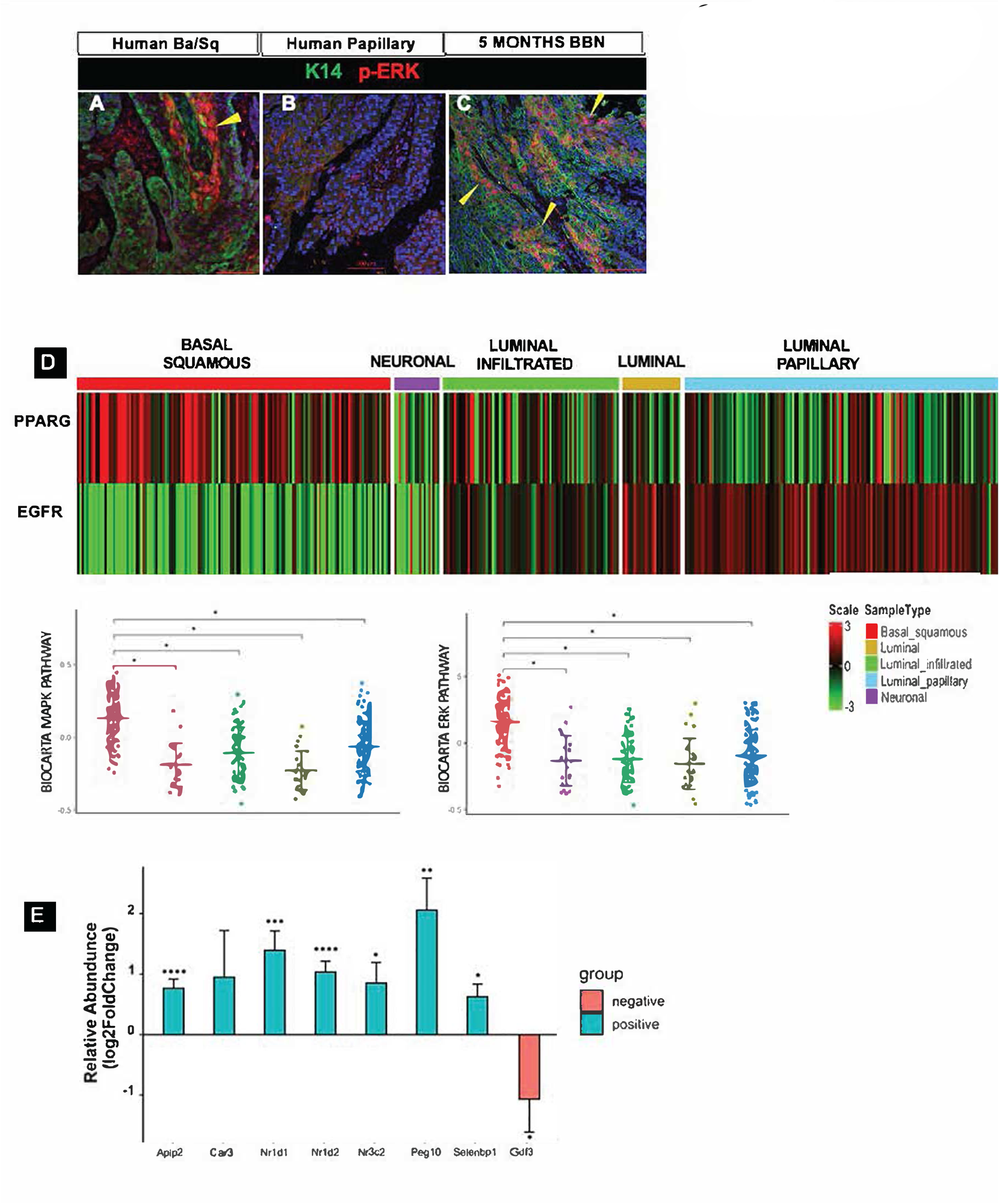
EGFR, MEK/ERK AND *PPARG* Expression and Pathways in Mouse and Human Tumors. **A.** p-ERK expression in a human Ba/Sq tumor. **B.** p-ERK expression in a human papillary tumor. **C.** p-ERK expression in a mouse BBN-induced Ba/Sq tumor. Bars=100μm. **D.** Expression of EGFR and *PPARG* and distribution of MAPK/ERK pathways among TCGA subtypes * p <0.01. **E.** Expression of the gene signature associated with dephosphphorylation of *Pparg* S273 residue after treatment with Rosi+Tram versus control in BBN 963 cells. Genes shown in green are up-regulated, genes shown in red are down-regulated.

**Supplementary Figure 1 related to Figure 3.**
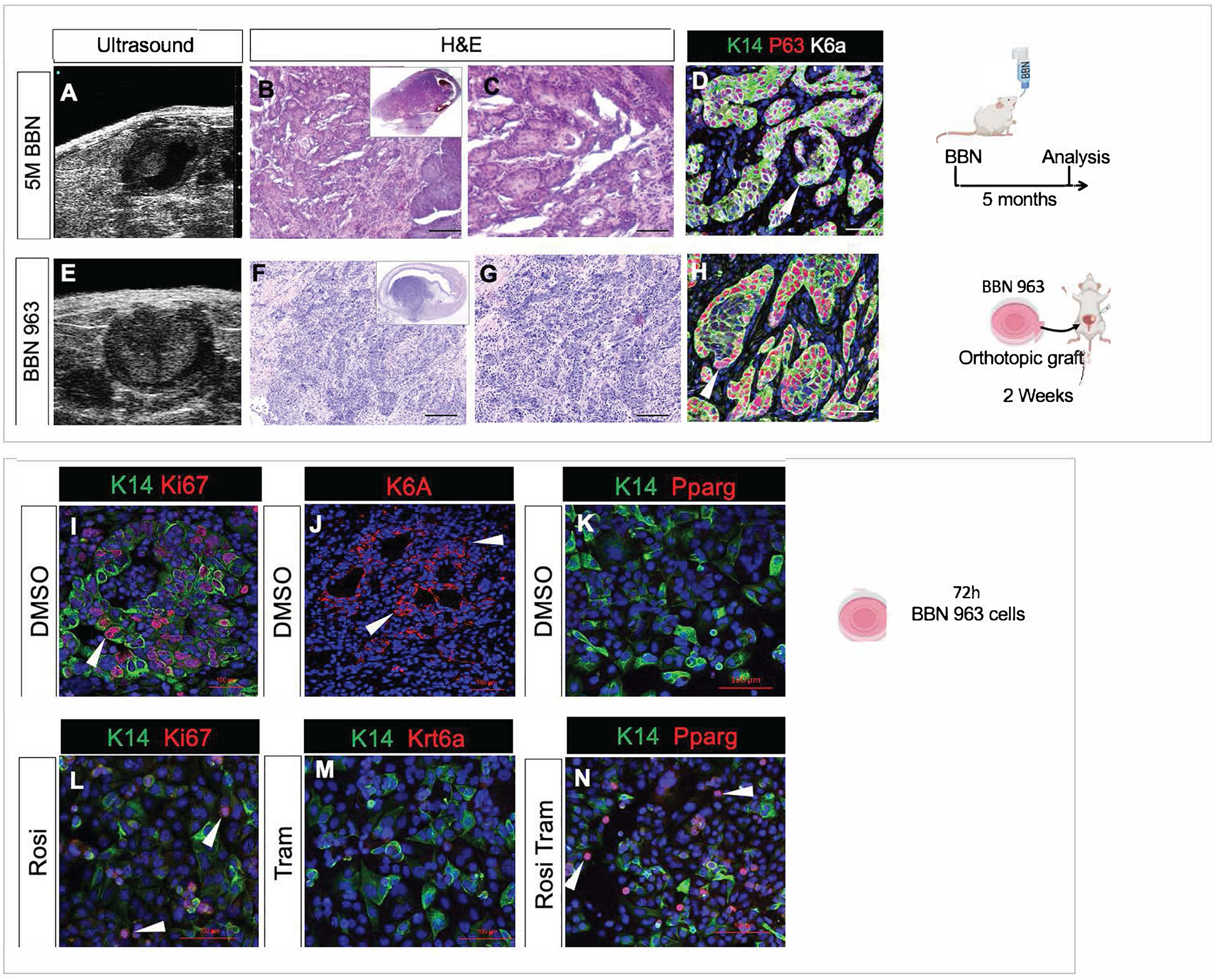
Comparison of the BBN 963 induced orthotopic tumors and BBN-induced tumors. **A-D.** Tumors from mice treated with BBN for 5 months analyzed by ultrasound (**A**), H/E staining (**B,C**) and expression of Ba/Sq markers, K14/P63/Krt6a (**D**). **E-H.** Characterization of orthotopic tumor grafts of BBN 963 cells by ultrasound (**E**), H/E staining (**F,G**) expression of Ki67, expression of Ba/Sq markers, K14/P63 and Krt6a (**H**). **I-N**. **Analysis of BBN 2-D cultures after 72h treatment with vehicle (I-K), Rosi (L), Tram (M) and Rosi+Tram.** I. BBN 963 cells treated with DMSO stained for Ki67. **J.** BBN 963 tumor cells treated with DMSO and Krt6a. **K.** K14/*Pparg* stained BBN 963 cells treated with DMSO. **L.** BBN 963 cells treated with Rosi stained for Ki67. **M.** BBN 963 tumor cells treated with Tram, stained for expression of Krt6a. **N.** BBN 963 cells treated with Rosi+Tram then stained for expression of *Pparg*. Scale bars: 50μm

**Supplementary Figure 2 Related to Figure 3.**
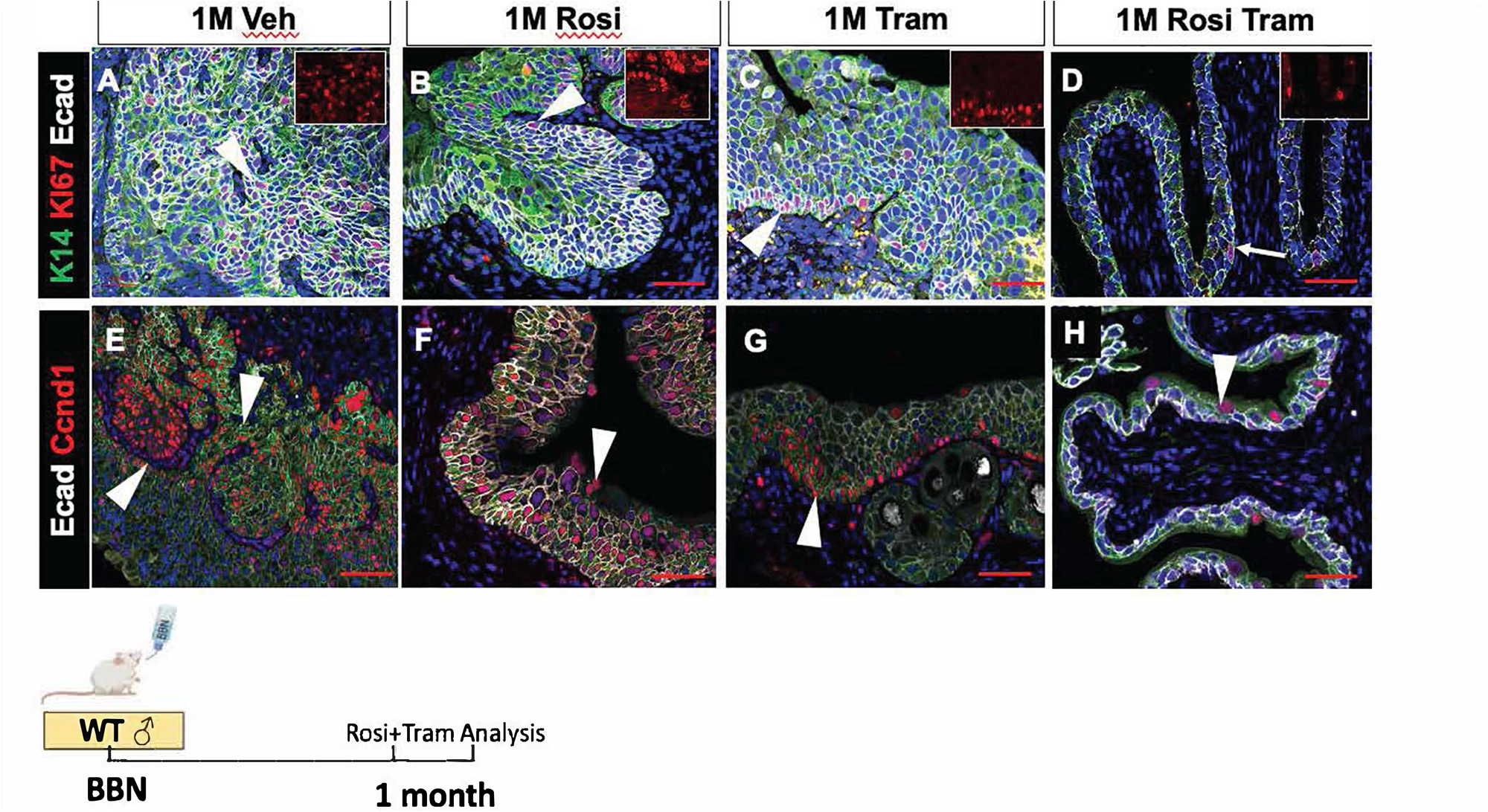
Expression of Ki67 and Ccnd1 in tumors from mice treated with BBN for 5 months, then with Rosi, Tram or Rosi+Tram for 1 month. **A-D.** Ki67 expression after 1 month of vehicle **(A)**, Rosi **(B)**, Tram **(C)** and Rosi+Tram **(D)**. **E-H.** Ccnd1 expression after 1 month of vehicle **(E)**, Rosi **(F)**, Tram **(G)** and Rosi+Tram **(H)**. Scale bars: 50μm

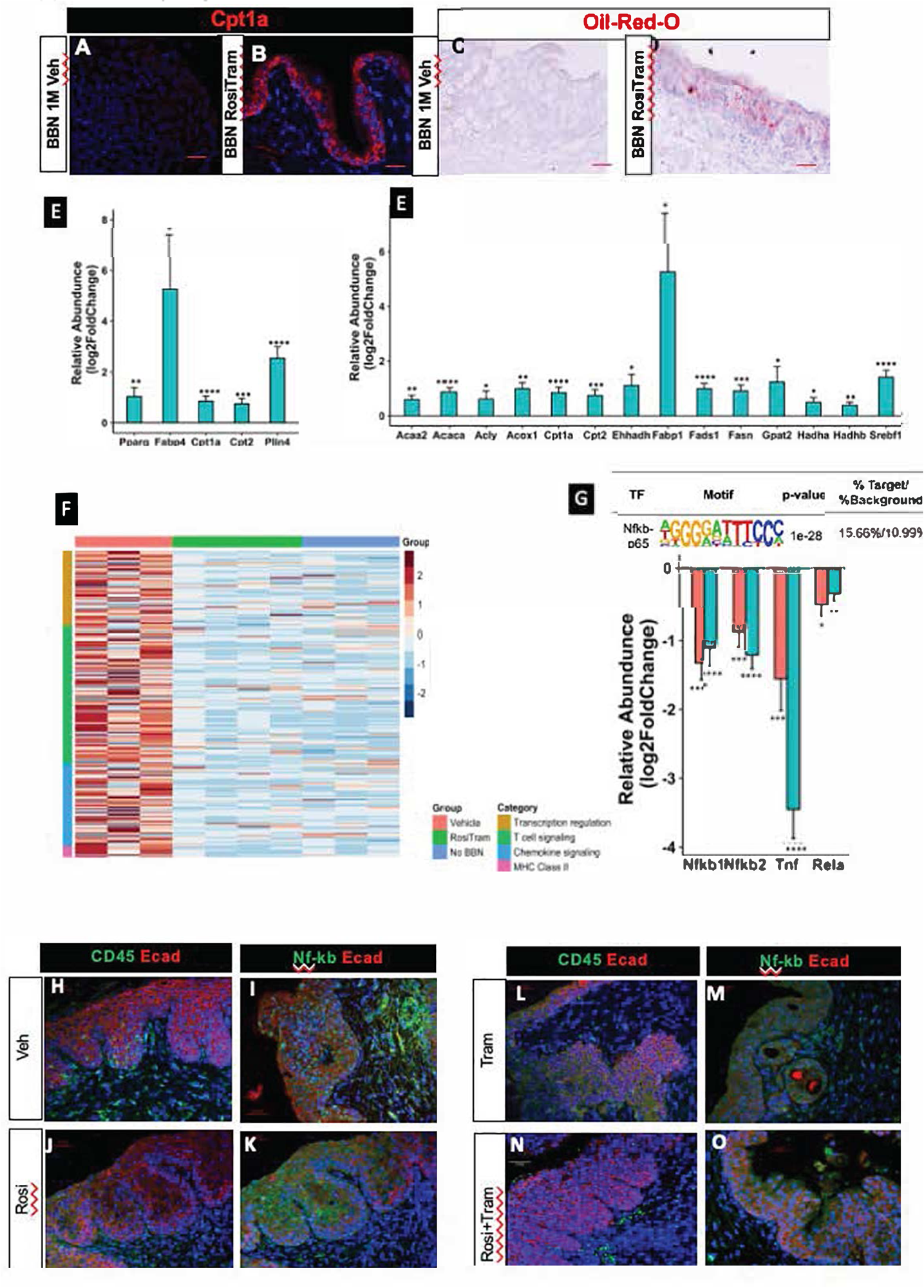

## Notes

### Competing Interest Statement

The authors have declared no competing interest.

